# Functional Robustness of Food Webs: A Dynamic Biomass Framework with Vital Species Sets and Cluster Influence

**DOI:** 10.64898/2026.08.27.747453

**Authors:** Xiaokang Qu, Chang Guo, Tianlong Fan, Linyuan Lü

## Abstract

1. Species loss can erode food-web functioning not only through secondary extinctions, but also through biomass redistribution, weakened energy pathways, and threshold-like functional collapse. Common topology-, connectivity-, and extinction-based robustness metrics provide valuable summaries of structural disassembly and cascade risk, but they are not designed to quantify continuous biomass retention, collapse-associated species sets, and non-additive group-level effects within a single dynamic framework.
2. We develop a dynamic biomass-based framework for assessing food-web robustness under progressive species removal. The framework introduces Dynamic Area-based Robustness (DAR), which quantifies the weighted area between slow- and fast-collapse reference trajectories of total ecosystem biomass retention. Building on these trajectories, we operationally define the Minimal Vital Species Set (MVSS) as the smallest fast-collapse-prefix species set whose removal first drives biomass below a predefined functional-collapse threshold. We further propose Cluster Influence (CI), which compares the biomass effect of simultaneous group removal with the mean effect of removing the same species individually.
3. We evaluated the framework using 120 niche-model virtual food webs spanning controlled gradients of species richness and connectance, and further demonstrated its applicability on 16 empirical stream food webs. We compared DAR with AUC- and secondary-extinction-based robustness metrics and assessed the sensitivity of DAR, MVSS, and CI to key bioenergetic parameters and parameter uncertainty.
4. DAR captured biomass-based robustness patterns that were only partly aligned with structural and extinction-based metrics, indicating that dynamic functional degradation provides complementary information. In virtual food webs, MVSS subsets were strongly enriched in basal species or basal resource nodes, and smaller MVSS proportions were associated with stronger positive CI under fast-collapse trajectories. Together, DAR, MVSS, and CI provide a reproducible framework for linking food-web structure, biomass dynamics, collapse thresholds, and non-additive species-set effects, offering a practical tool for dynamic robustness assessment in theoretical and empirical food webs.

## 1 Introduction

Species loss can alter food-web functioning not only by causing extinctions, but also by redistributing biomass, weakening energy pathways, and pushing ecosystems toward functional collapse. Anthropogenic disturbance, biological invasions, and climate change can therefore trigger cascading effects that compromise ecosystem stability and functioning (Wilmers, 2007). Food webs provide a natural framework for evaluating such responses because they represent trophic dependencies and the pathways through which energy and biomass are transferred among species.

A large body of research has assessed food-web robustness using static topological metrics, including connectance, modularity, nestedness, and related structural indices (Borrvall et al., 2000; Estrada, 2006; Rooney and McCann, 2012; Garay-Narváez et al., 2014; Abernethy et al., 2019; Keyes et al., 2024). These metrics are interpretable, computationally efficient, and useful for comparing broad network architectures. However, when calculated from presence/absence topology alone, these metrics treat species as nodes and interactions as links, and therefore cannot by themselves represent how biomass redistributes after species loss, how compensatory dynamics attenuate or amplify perturbations, or when energy flow falls below a functionally critical level.

Secondary-extinction (SE) frameworks evaluate extinction cascades triggered by primary species removals. In their simplest form, these simulations are implemented on qualitative food-web networks, where consumers are removed when they lose all trophic resources (Paine, 1966; Pimm, 1980; Sole and Montoya, 2001). Such approaches have revealed important patterns, including robustness to random species loss and fragility to targeted removals (Dunne et al., 2004; Dunne and Williams, 2009; Ebenman et al., 2004; Ståhl et al., 2025), and have also been applied to mutualistic systems where network architecture can promote tolerance to disturbance (Memmott et al., 2004). More refined SE simulations can incorporate weighted interactions, extinction thresholds, or quantitative food-web information, thereby relaxing the assumption that consumers persist until all resources are lost (Bellingeri and Vincenzi, 2013). These developments show that SE simulations can be formulated at different levels of biological detail, from qualitative cascade models to quantitative implementations.

Another class of robustness metrics summarizes structural disassembly using area-under-the-curve (AUC)-based measures. In this approach, species are sequentially removed according to a specified removal sequence, the size of the largest connected component (LCC) is recorded after each removal step, and the area under the resulting LCC-retention curve is used as a scalar measure of structural robustness (Albert et al., 2000; Schneider et al., 2011). Depending on the data available, AUC-based robustness metrics can be calculated on qualitative food-web networks or adapted to quantitative networks that include interaction strengths, energy fluxes, or temporal variation (Kortsch et al., 2021). In common presence/absence applications, these metrics describe how rapidly the connected component declines during species removals, thereby providing useful summaries of structural robustness (Gao et al., 2016). However, when the AUC is calculated only from connectivity-based LCC trajectories, it should be interpreted as a measure of structural disassembly rather than a direct measure of biomass production, energy transfer, or functional viability. Additional quantitative or dynamic information is therefore required to evaluate whether ecosystem functioning has degraded beyond a biologically meaningful threshold.

A related line of work seeks to identify important species or groups of species. Eigenvector-based approaches, for example, rank species according to their contribution to co-extinction risk and can identify species whose loss disproportionately accelerates network collapse (Allesina and Pascual, 2009). Other approaches have considered the idea of keystone species complexes, where multiple species jointly contribute to ecosystem structure or functioning (Ortiz et al., 2017). These studies are important because ecological stability may depend not only on single keystone species, but also on combinations of species whose collective roles preserve critical energy pathways. However, many existing approaches primarily rank species or identify influential groups from network structure or interaction patterns, rather than defining the retained species set required to keep a dynamic ecosystem function above a collapse threshold.

Recent advances have also incorporated rewiring, extinction propagation, quantitative interaction information, and dynamic food-web responses. Rewiring-based models evaluate how adaptive changes in trophic interactions can buffer extinction cascades and alter robustness estimates (Staniczenko et al., 2010; Ávila-Thieme et al., 2023). Quantitative and biomass-informed food-web studies have further shown that ecosystem responses to disturbance depend on interaction strengths, energy pathways, and the distribution of biomass or flux among species (Lewis et al., 2022; Rodriguez and Saravia, 2024). Dynamic deletion models go further by simulating biomass responses during sequential species removals, showing that robustness estimates can differ substantially between static and dynamic food webs (Brose et al., 2006; Curtsdotter et al., 2011; Delmas et al., 2017). Together, these studies demonstrate the importance of trophic traits, energetic constraints, biomass feedbacks, and quantitative food-web structure. However, to our knowledge, few existing frameworks simultaneously quantify continuous functional degradation, apply an explicit functional-collapse threshold, identify collapse-preventing retained species sets, and measure non-additive group-level effects.

These limitations motivate three methodological needs. First, robustness should be quantified from biomass trajectories rather than inferred only from network structure, connectivity loss, or extinction counts. Second, robustness assessment should identify not only individually important species, but also retained species sets that keep ecosystem function above a collapse threshold. Third, because species can interact non-additively, robustness analysis should quantify whether groups of species have synergistic, redundant, or compensatory effects on biomass loss. Existing metrics address important components of these problems, but they rarely address all three within a single framework.

Building on this literature, we develop a dynamic, biomass-based framework for food-web robustness grounded in species-loss simulations. First, we introduce Dynamic Area-based Robustness (DAR), which quantifies functional degradation from simulated biomass trajectories relative to an explicit collapse threshold. Second, we identify the Minimal Vital Species Set (MVSS), defined here as the species set found under our greedy search procedure whose persistence keeps total biomass above a predefined functional-collapse threshold. MVSS is related to the idea of a keystone species complex (Ortiz et al., 2017), but is operationally defined through dynamic species-loss simulations and an explicit biomass-collapse criterion. Third, we propose the Cluster Influence (CI) metric, which quantifies group-level interaction effects by comparing the biomass consequences of simultaneous group removals with the mean consequences of removing the same species individually. Together, DAR, MVSS, and CI provide a unified framework for evaluating dynamic robustness, identifying MVSSs, and measuring non-additive group effects in food webs.

We evaluate DAR, MVSS, and CI using 120 virtual food webs generated with the niche model (Williams and Martinez, 2000), spanning systematically varied species richness and connectance, and further demonstrate the framework on 16 empirical stream food webs. We then assess the robustness of our conclusions through parameter-sensitivity analyses of key bioenergetic parameters and uncertainty quantification based on Latin Hypercube Sampling. Finally, we compare DAR directly with AUC-and SE-based metrics to determine what complementary information dynamic biomass-based robustness provides relative to existing connectivity-, AUC-, and extinction-cascade-based measures. Table 1 summarises how the proposed DAR–MVSS–CI framework compares with existing robustness metrics, including the ecological questions each metric addresses, its main strengths, and its limitations relative to dynamic biomass-based robustness assessment.

**Table 1:** Comparison of existing food-web robustness metrics with the DAR–MVSS–CI framework.

| Metric | Reference(s) | Question addressed | Response | Collapse thr. | Critical set | Group effects | Main strength | Limitation relative to DAR-MVSS-CI |
| --- | --- | --- | --- | --- | --- | --- | --- | --- |
| Connectance | Dunne et al. (2002); Gilbert (2009); Abernethy et al. (2019); Keyes et al. (2024) | How does interaction density affect robustness to species loss? | Topology | ✗ | ✗ | ✗ | Simple and widely used descriptor of interaction density linked to structural robustness. | Does not by itself model biomass redistribution, functional collapse thresholds, MVSS, or group effects. |
| Modularity | Rooney and McCann (2012); Garay-Narváez et al. (2014); Keyes et al. (2024) | How does compartmentalised food-web structure affect disturbance spread and species persistence? | Topology / compartment structure | ✗ | ✗ | ✗ | Captures compartment structure that can buffer perturbation spread in some contexts. | Mostly structural; effects are context-dependent and do not identify biomass-collapse thresholds, MVSS, or group synergy. |
| Nestedness | Memmott et al. (2004); Keyes et al. (2024) | How does nested interaction structure affect tolerance to species loss? | Topology | ✗ | ✗ | ✗ | Describes ordered interaction structure and potential extinction tolerance. | Primarily structural; evidence is strongest in mutualistic networks and less central in food-web robustness studies. |
| AUC | Albert et al. (2000); Schneider et al. (2011); Kortsch et al. (2021) | How does a selected network-level response change during sequential species removal? | Connectivity / selected network response | ✗ | ✗ | ✗ | Provides a scalar summary of robustness across removal sequences and can be applied to qualitative or quantitative network responses. | When calculated only from connectivity, it measures structural disassembly rather than biomass-based functional degradation or collapse thresholds. |
| SE | Paine (1966); Pimm (1980); Sole and Montoya (2001); Dunne et al. (2004); Dunne and Williams (2009); Ebenman et al. (2004) | How many additional species are lost after primary removals trigger extinction cascades? | Extinction cascade | ✗ | ✗ | ✗ | Captures extinction cascades and is widely applicable to ecological networks. | Classic qualitative implementations use discrete extinction criteria; SE alone does not identify MVSS or quantify non-additive biomass effects. |
| Weighted / threshold-based robustness | Bellingeri and Vincenzi (2013); Ståhl et al. (2025) | How do interaction strengths or extinction thresholds alter robustness estimates? | Weighted / threshold-based extinction cascade | ✓ | ✗ | ✗ | Adds graded extinction criteria beyond simple loss of all trophic resources. | Does not generally integrate continuous biomass-retention trajectories with MVSS identification and CI-based group-effect assessment. |
| Eigenvector co-extinction ranking | Allesina and Pascual (2009) | Which species are most important for accelerating co-extinction-driven collapse? | Co-extinction ranking | ✗ | ✗ | ✗ | Efficiently ranks species using full-network topology rather than local degree alone. | Ranks individual species and collapse sequences, but does not identify retained species sets that prevent biomass-based functional collapse. |
| Keystone species complex | Ortiz et al. (2017) | Which groups of species jointly contribute to ecosystem structure or functioning? | Species-group importance | ✗ | ✗ | ✗ | Recognises that ecosystem functioning can depend on groups of species rather than only single keystone species. | Identifies important species complexes, but not through dynamic biomass-collapse simulations or an explicit retained-set threshold such as MVSS. |
| Rewiring-based robustness | Staniczenko et al. (2010); Ávila-Thieme et al. (2023) | How does adaptive rewiring affect extinction propagation after species loss? | Rewiring / extinction propagation | ◦ | ✗ | ✗ | Captures compensatory changes in trophic interactions after perturbation. | Focuses on extinction propagation and rewiring rather than biomass-based collapse or group synergy. |
| Bioenergetic dynamic models | Brose et al. (2006); Curtsdotter et al. (2011); Delmas et al. (2017) | How do biomass dynamics and trophic traits affect food-web persistence under perturbation or deletion? | Biomass trajectory | ◦ | ✗ | ✗ | Represents energetic constraints, biomass feedbacks, and compensatory responses. | May quantify dynamic persistence or extinction thresholds, but does not formally identify MVSS or quantify non-additive group effects. |
| DAR + MVSS + CI | **This study** | How can dynamic robustness, MVSSs, and non-additive group effects be quantified within one framework? | Biomass trajectory | ✓ | ✓ | ✓ | Integrates dynamic robustness, MVSS, and CI within one biomass-based framework. | — |

*Notes:* Response indicates the main aspect of network response tracked by each metric. Collapse threshold indicates whether the method uses an explicit threshold for functional or extinction-related collapse. Critical species set indicates whether the method identifies a retained species set sufficient to keep ecosystem function above an explicit collapse threshold. Group effects indicate whether non-additive multi-species effects are quantified. ✓ = yes; ✗ = no; ◦ = partial.

## 2 Materials and methods

### 2.1 Food webs

We used two complementary food-web datasets. The primary analyses were conducted on virtual food webs generated under controlled structural conditions, whereas the empirical analyses were retained as a secondary demonstration of the framework on realistic stream food webs.

Virtual food webs were generated using the niche model (Williams and Martinez, 2000), a widely used model for generating food-web structures from specified species richness and connectance. Species richness was set to *S* = 20, 50, and 100, and connectance was set to *C* = 0.05, 0.10, 0.15, and 0.20. For each *S × C* combination, we generated 10 replicate food webs, yielding 120 virtual food webs in total. This factorial design allowed us to evaluate DAR, MVSS, and CI across controlled gradients of network size and interaction density before applying the framework to empirical food webs.

We additionally applied the framework to 16 empirical stream food webs from New Zealand streams spanning different surrounding catchment (Thompson and Townsend, 1999; Thompson et al., 2001; Thompson and Townsend, 2003, 2005). These are aquatic stream food webs; the habitat labels reported in Table S2 refer to the surrounding catchment vegetation or land-use context rather than to terrestrial food webs. The empirical food webs were used to demonstrate applicability to realistic food-web topologies rather than as the primary basis for inference. They ranged from 49 to 113 food-web nodes, which we refer to as species hereafter for consistency with the model terminology, and from 0.0298 to 0.0655 connectance. Full structural properties of the empirical stream food webs are reported in Table S2. We selected these 16 stream food webs because they form a coherent empirical dataset with comparable sampling protocols, binary trophic matrices, and variation in species richness, connectance, and catchment context, allowing the framework to be demonstrated on realistic but methodologically comparable food-web topologies.

Both virtual and empirical food webs were represented as binary directed adjacency matrices, where 0 and 1 indicate the absence and presence of trophic links, respectively. Links were treated as directed trophic interactions. Before analysis, all matrices were standardized by removing duplicate links and self-loops.

For both virtual and empirical food webs, we quantified species richness (*S*) and connectance (*C*), because these two quantities define the controlled virtual-web design and provide the most direct comparison between virtual and empirical food webs. For empirical food webs, we additionally reported modularity (*M*) and global efficiency (*GE*) as descriptive structural summaries of network compartmentalisation and overall structural integration.

### 2.2 Trophic-level calculation

To characterize the trophic composition of whole food webs and MVSS subsets, we used two complementary classifications. First, each species was assigned to a topological trophic role based on its position in the directed food-web matrix. Species without prey were classified as producers, species with both prey and predators were classified as intermediate consumers, and species without predators were classified as top predators. This three-class role classification was used for the main trophic-role composition analysis.

Second, we calculated continuous (prey-averaged) trophic levels following the standard formulation (Levine, 1980; Williams and Martinez, 2004). Let *A* denote the directed predator–prey adjacency matrix, where *A_ij_* = 1 indicates that species *i* consumes species *j*. For each consumer *i*, diet proportions were calculated as

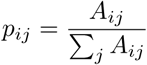

with basal species (those without prey) assigned trophic level 1 (equivalently *p_ij_* = 0). Trophic levels were then obtained by solving

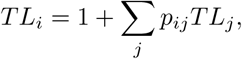

or equivalently the linear system (*I − P*) **TL** = **1** for all species simultaneously, where *P* = [*p_ij_*]. Self-loops (cannibalistic links) were removed during preprocessing prior to this calculation. For visualization, continuous trophic levels were grouped into four broad trophic classes: basal producers (*TL* = 1), low-level consumers (1 *< TL ≤* 2), mid-level consumers (2 *< TL ≤* 3), and higher-level consumers (*TL >* 3). The same trophic-level calculation and broad-class grouping were used for the virtual and empirical trophic-composition summaries.

### 2.3 Bioenergetic dynamic model

To capture biomass-mediated species interactions, we used an allometric bioenergetic model of food-web dynamics. This class of models has been widely used to simulate consumer–resource biomass dynamics, energetic constraints, metabolic losses, and extinction responses in complex food webs (Yodzis and Innes, 1992; Brose et al., 2006; Williams et al., 2007; Delmas et al., 2017). The model describes temporal changes in species biomass and is given in Equations (1a) and (1b).

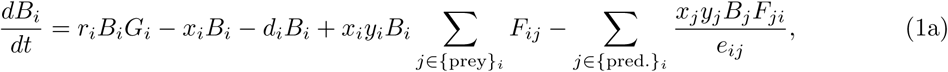

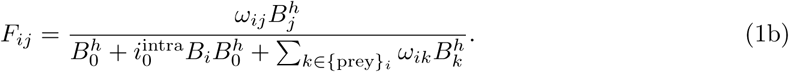

The bioenergetic model (Eq. (1a)) describes the temporal dynamics of species biomass *B_i_* within a food web, incorporating primary production, metabolic loss, natural mortality, predation gains, and losses. Specifically, the rate of change in biomass of species *i* is governed by:

- **Primary production**: for basal species, the intrinsic growth term of *i* is represented by *r_i_B_i_G_i_*, where *r_i_* is the intrinsic growth rate and *G_i_* is the logistic net growth rate with *G_i_* = 1 *−* (*B_i_/K_i_*), *K_i_* being the carrying capacity of *i*.
- **Body mass**: species body masses were assigned according to trophic position using a constant consumer–resource body-mass ratio *Z*, such that *M_i_* = *Z ^T^ ^Li^*^−1^, where *TL_i_* is the continuous (prey-averaged) trophic level of species *i* defined in Section 2.2 (basal species have *TL_i_* = 1 and therefore *M_i_* = 1). Body mass thus increases with trophic level, and all metabolic, consumption, and mortality rates scale allometrically with *M_i_*. Unless otherwise stated, we used a default ratio of *Z* = 10.
- **Metabolic loss**: species expend biomass at a metabolic rate *x_i_*, resulting in the loss term *−x_i_B_i_*.
- **Natural mortality**: the natural mortality rate *d_i_* contributes a further loss *−d_i_B_i_*.
- **Predation gain**: for consumer *i*, biomass gain is modelled by

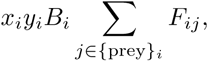

where *y_i_* is the maximum consumption rate relative to its metabolic rate. *F_ij_* (Eq. (1b)) is the multi-resource functional response of *i* consuming *j*. *ω_ij_* denotes the preference of predator *i* for prey *j* (*ω_ij_* = 1*/|{*prey*}_i_|* by default, assuming uniform preference). *B*_0_ is the half-saturation constant that controls how feeding rates saturate with prey biomass. *i*^intra^ is the intensity of intraspecific interference among consumers, and *h* is the Hill exponent that shapes the functional response: if *h* = 1 and *i*^intra^ = 0, *F_ij_* reduces to a Holling type II response; if *h >* 1 and *i*^intra^ = 0, it becomes a Holling type III response; if *i*^intra^ *>* 0, interference-modified forms are recovered, dampening predation efficiency.

- **Predation loss**: species *i* may be preyed upon by predators *j ∈ {*pred.*}_i_*, resulting in the loss term

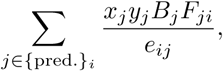

where *e_ij_*is the assimilation efficiency of predator *i* consuming prey *j* (so that, in the predation-loss term, *e_ij_* applies to predator *j* feeding on species *i*).

The model allows flexible modelling of both basal and consumer species, accommodates different feeding strategies, and reflects realistic saturation and interference effects in consumer-resource interactions. Default parameter settings are listed in Table S1.

### 2.4 Evaluation framework

We developed an integrated framework to evaluate food-web robustness under biomass-dynamic conditions. Total ecosystem biomass (TEB) was used as a proxy for ecosystem functioning, and species-level biomass trajectories were simulated with the bioenergetic dynamic model described above. The framework links three complementary quantities: Dynamic Area-based Robustness (DAR), Minimal Vital Species Set (MVSS), and Cluster Influence (CI).

DAR quantifies the range of biomass-retention outcomes generated by different removal sequences, where each sequence specifies an order in which species are sequentially deleted from the same food web. MVSS identifies the species set, obtained as the smallest prefix of the FCT whose removal first drives TEB below a predefined collapse threshold, such that the persistence of these species prevents that collapse event. CI quantifies the non-additive influence of species groups on biomass dynamics and is used to evaluate whether multi-species effects help explain the functional importance of MVSSs. The following subsections describe the calculation of DAR, MVSS, CI, and the structural comparison metrics, respectively.

#### 2.4.1 Dynamic Area-based Robustness (DAR)

### Definition and motivation

DAR quantifies the resistance of food-web functioning to progressive species loss under biomass-dynamic conditions. We used TEB as a proxy for ecosystem functioning, because aggregate biomass is widely used in biodiversity–ecosystem functioning research to represent ecosystem-level production and functioning (Tilman et al., 2006; Hooper et al., 2005; Maureaud et al., 2019).

Let *B*^(0)^ = (*B*^(0)^_1_, …, *B*^(0)^_*S*_) denote the initial biomass vector of the *S* species, and let *B*^∗^ = (*B*^∗^*, . . ., B*^∗^) denote the equilibrium biomass vector. Then

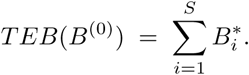

During progressive species loss, species are removed one at a time according to a removal sequence, defined here as the ordered list of species deleted during the simulation. After each deletion, the bioenergetic model is re-simulated to equilibrium. For a removal set *K ⊆* 1*, . . ., S*, the resulting equilibrium biomass vector is denoted *B*^(−^*^K^*^)^, and the corresponding TEB is

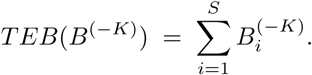

The trajectory of TEB across removal steps therefore provides a quantitative description of how food-web functioning changes as extinctions accumulate.

### Perturbations to ecosystem functioning

In a given food web, defined by its species pool and trophic interactions, ecosystem functioning can be altered by species loss or gain, shifts in interaction strengths, or changes in species-specific traits such as metabolic rate. Here, we focused on sequential species-removal simulations because they are widely used in food-web robustness analyses to represent progressive biodiversity loss and extinction cascades (Dunne et al., 2004; Dunne and Williams, 2009; Curtsdotter et al., 2011; Delmas et al., 2017). By applying a removal sequence and recording TEB after each removal step, we obtained a biomass-retention trajectory that characterizes the food web’s dynamic response to progressive species loss. This response is not necessarily monotonic: removing a species may increase or decrease TEB depending on its trophic position, functional role, and indirect effects.

### Heuristic patterns from empirical studies

Empirical and theoretical studies support general heuristic rules for simple food webs. The removal of top predators, which release lower trophic levels from suppression, or the elimination of redundant competitors, which overlap strongly in niche and provide little unique function, can sometimes increase TEB (Pace et al., 1999; Schmitz et al., 2000). In contrast, the loss of key producers that dominate primary production, or of species functioning as energy hubs that connect multiple trophic levels, often leads to sharp biomass declines (Hooper et al., 2005; Ives and Carpenter, 2007). However, in more complex food webs—characterized by high connectance or broad trophic generality—indirect interactions such as omnivory and apparent competition can override these patterns, making them unreliable (Bascompte et al., 2005; Gellner and McCann, 2012).

### Construction of slowest-and fastest-collapse trajectories

Exhaustively evaluating all possible removal sequences is computationally infeasible for food webs with many species. We therefore used a greedy heuristic to generate two limiting biomass-retention trajectories. At each step, the SCT removes the species whose deletion leaves the highest remaining TEB after re-simulation, thereby minimizing immediate biomass loss. The FCT is defined as the reverse of the SCT, providing a mirrored heuristic trajectory that drives the system toward collapse more rapidly.

By construction, the FCT is the exact reverse of the SCT. This symmetric design allows species removed early in the FCT to be compared with species retained until late in the SCT. Although these trajectories are heuristic rather than globally optimal, they provide a tractable and comparable basis for bounding food-web responses under progressive species loss.

Together with a predefined collapse threshold, the SCT and FCT define the feasible robustness range: the set of removal–biomass states for which the food web avoids functional collapse. The area enclosed by the trajectories and the collapse threshold quantifies resistance to collapse, with larger areas indicating greater robustness and smaller areas indicating greater fragility.

Algorithm 1 formalizes the construction of the SCT and FCT, using three inputs: the parameterized bioenergetic model, the initial biomass vector, and the simulation horizon (*t* = 500). The horizon was validated through sensitivity tests (Figure S1), and initial species biomass was randomly assigned within (0, 1).

Two critical limitations of this algorithm should be emphasized:

- the SCT is a **local heuristic trajectory**, not a guaranteed globally optimal slowest-collapse sequence;
- the FCT is a **mirrored heuristic trajectory**, not a guaranteed globally optimal fastest-collapse sequence.

**Algorithm 1.**
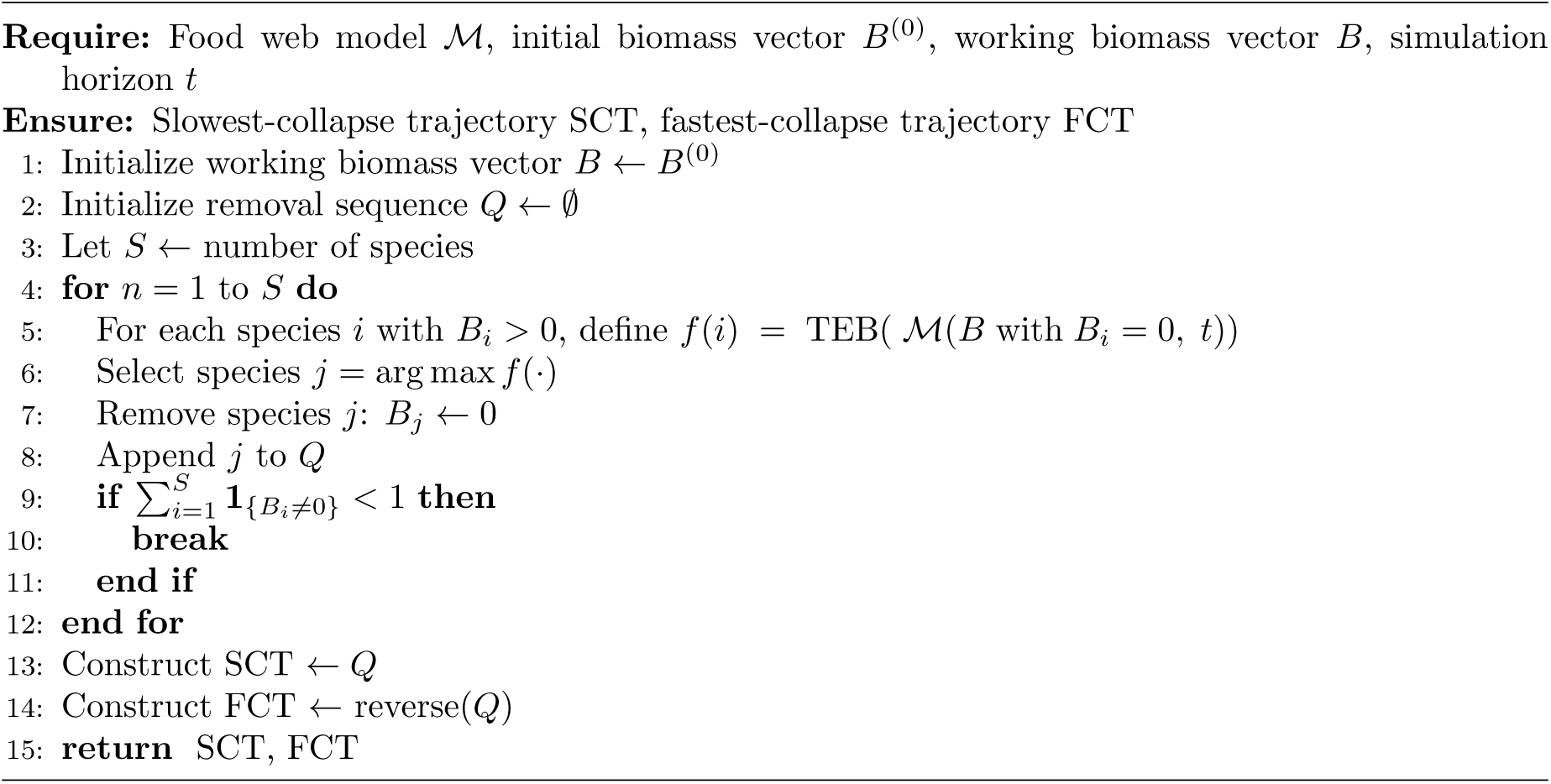
Greedy algorithm to generate the SCT and the FCT.

### Collapse threshold and DAR

We assessed food-web robustness using the SCT and the FCT, together with a functional-collapse threshold motivated by the IUCN Red List of Ecosystems protocol (Bland et al., 2017), which evaluates ecosystem risk in relation to functional collapse. In this study, a food web was considered functionally collapsed when its TEB fell below 10% of its initial value. We therefore define the relative collapse threshold as *θ* = 0.1, and the corresponding absolute TEB threshold as

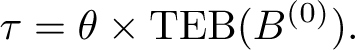

The relative threshold *θ* is used when calculating biomass-retention curves and DAR, whereas the absolute threshold *τ* is used to determine whether a simulated food web has crossed the functional-collapse threshold.

Figure 1a illustrates the calculation of DAR. The *x*-axis represents the fraction of species removed relative to the total number of species, and the *y*-axis represents the fraction of TEB retained relative to the initial TEB. The FCT approximates a fast-collapse trajectory, whereas the SCT approximates a slow-collapse trajectory. The collapse threshold is therefore marked at the relative value *θ* = 0.1.

**Figure 1:**
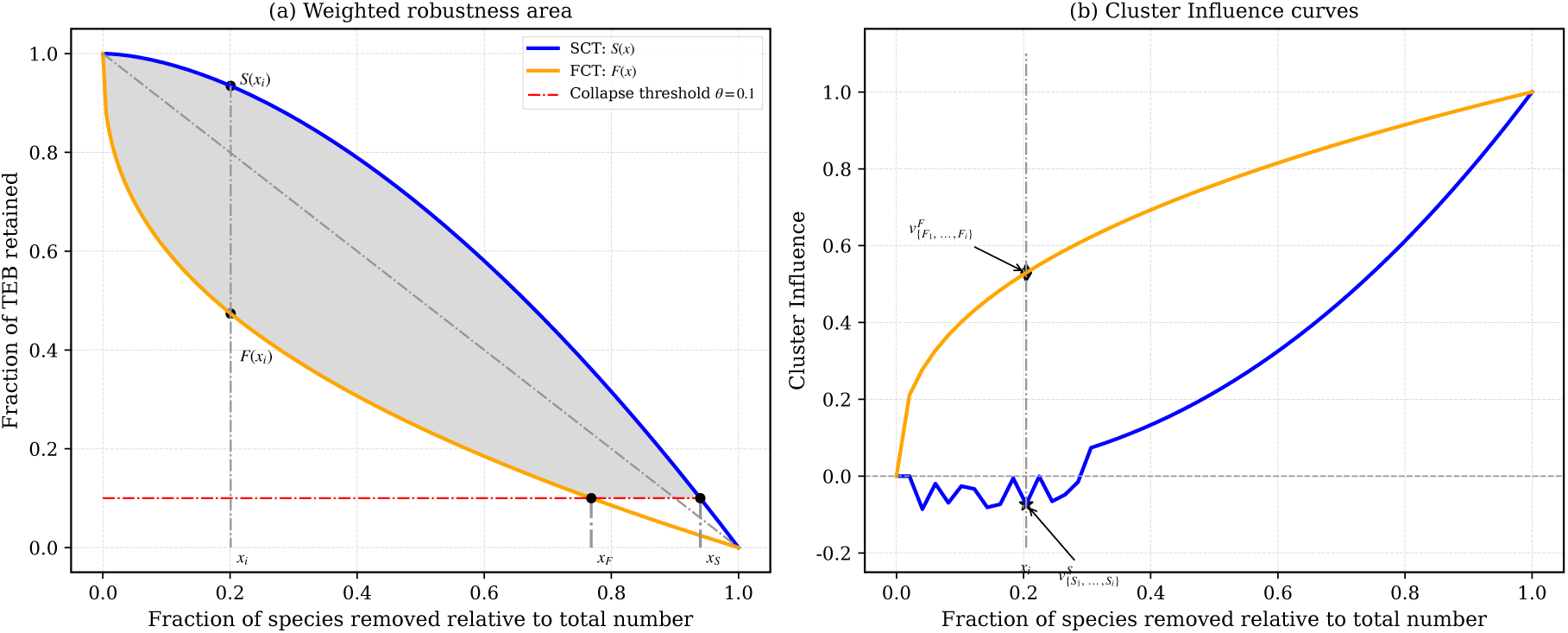
Illustrative examples of the DAR–MVSS–CI framework. (a) Weighted robustness area. The shaded region represents the food web’s resistance to functional collapse under progressive species removal. The area integrates biomass-retention ratios between the SCT, the FCT, and the relative collapse threshold *θ* = 0.1, with early removals weighted more heavily. (b) Cluster Influence curves. The curves show *v^F^* (*x*) and *v^S^*(*x*), the Cluster Influence values for prefix species sets along the FCT and SCT, respectively, as the fraction of removed species increases.

To formalize the robustness area, let *S*(*x*) and *F* (*x*) denote the TEB retention ratios along the SCT and FCT, respectively, at a given removal fraction *x*: *S*(*x*) = TEB_SCT_(*x*)*/*TEB(*B*^(0)^) and *F* (*x*) = TEB_FCT_(*x*)*/*TEB(*B*^(0)^). Let *x_F_* be the first removal fraction at which the FCT reaches the relative collapse threshold, and let *x_S_* be the first removal fraction at which the SCT reaches the relative collapse threshold. Thus, *F* (*x_F_*) = *θ* and *S*(*x_S_*) = *θ*.

DAR is then defined as the sum of two weighted integrals. The first integral, from *x* = 0 to *x* = *x_F_*, is the weighted area between the SCT and FCT before the FCT reaches collapse. The second integral, from *x* = *x_F_* to *x* = *x_S_*, is the weighted area between the SCT and the relative collapse threshold after the FCT has already collapsed:

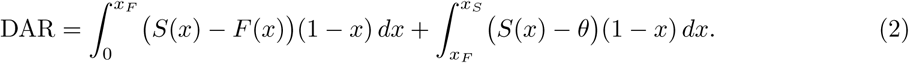

In practice, species removal is a discrete process because one species is removed at each step. We therefore computed the weighted integrals numerically using a summation with step size Δ*x* = 1*/S*, where *S* is the number of species in the food web. For discrete step *i* (*i* = 0, 1*, . . ., S*), the robustness area was approximated as

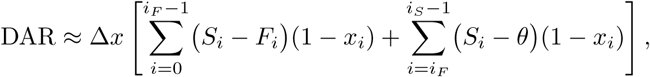

where *i_F_*and *i_S_* are the first discrete steps at which the FCT and SCT reach the relative collapse threshold *θ*, respectively, and *S_i_*and *F_i_* are the corresponding TEB retention ratios at step *i*.

### Rationale for weighting

The factor (1 *−x*) gives greater weight to biomass loss occurring at earlier stages of species removal. This reflects the assumption that biomass loss in a relatively intact food web indicates a stronger reduction in resistance to progressive species loss than the same absolute loss occurring after most species have already been removed. At *x ≈* 0, a given biomass reduction occurs when the food web is still largely intact; at *x ≈* 1, the same absolute reduction occurs when collapse has already progressed. Therefore, (1 *− x*) down-weights late-stage removals and emphasizes early-stage functional degradation, ensuring that DAR reflects vulnerability to early biodiversity loss under progressive disturbance.

#### 2.4.2 Minimal Vital Species Set (MVSS)

The SCT and the FCT characterize two limiting biomass-retention trajectories of a food web under progressive species loss (Section 2.4.1). While these trajectories summarize system-level degradation, they do not by themselves identify which species collectively form the critical set associated with functional collapse. We therefore define the MVSS as the smallest FCT prefix whose removal first causes TEB to fall below the collapse threshold. In this operational sense, MVSS identifies a collapse-associated vital species set: the removal of this set is sufficient to trigger the collapse event along the FCT, and the continued presence of these species prevents that specific FCT-defined collapse event. This definition should not be interpreted as a globally minimal retained set under all possible removal combinations.

### Definitions

To ensure consistency with the dynamic simulation framework, we used the following operational definitions:

- **Collapse threshold:** A food web is considered collapsed when its *TEB* satisfies

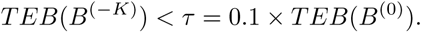

- **Species removal:** Removing species *i* corresponds to setting its initial biomass to zero before simulation, *B_i_*^(0)^ = 0.
- **Species presence:** A species was considered present, or viable, if its equilibrium biomass satisfied *B_i_^*^* > 10^−4^ and its biomass change rate was below 10^−4^.
- **MVSS:** The MVSS was defined as the smallest prefix of the FCT whose removal first causes functional collapse.

### Identification of MVSS from the FCT

The MVSS was obtained directly from the FCT using a reproducible two-step procedure.

- **Step 1: Locate the collapse index on the FCT.** Let (*ϕ*_1_*, ϕ*_2_*, . . ., ϕ_S_*) denote the ordered species removals along the FCT. We computed

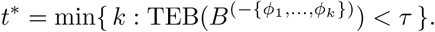

where *t*^∗^ is the first removal step that induces functional collapse. Removing *ϕ*_1_*, . . ., ϕ_t_∗*_−1_ keeps the food web above the collapse threshold, whereas removing *ϕ_t_∗* causes collapse.

- **Step 2: Construct the MVSS.** The MVSS was defined as the FCT prefix up to and including the first collapse-inducing removal:

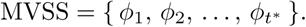

Species *ϕ_t_∗* was included because its removal is the critical event that first pushes TEB below the collapse threshold. Thus, the persistence of the species in the MVSS, together with their trophic links, is sufficient to prevent the collapse event identified along the FCT.

This definition depends only on the simulated biomass response under progressive species removal and can therefore be applied consistently across virtual and empirical food webs.

### Methodological remarks

The MVSS reflects a vital species set derived from the heuristic construction of the FCT. Because the FCT is generated from a greedy procedure rather than an exhaustive search over all possible species subsets, the MVSS should be interpreted as an algorithmically identified collapse-associated vital set, not as a guaranteed globally minimal species set. Alternative species sets of comparable or smaller size may exist, but the proposed procedure provides a tractable and reproducible approximation for comparing food webs.

#### 2.4.3 Cluster Influence (CI)

Species can exert non-additive effects on ecosystem functioning: the combined loss of multiple species can produce impacts that are larger, smaller, or qualitatively different from expectations based on individual species effects (Sih et al., 1998; Loreau and Hector, 2001; Ball et al., 2008; Liu et al., 2020; Guo et al., 2025). To quantify such group-level effects under biomass dynamics, we define the CI metric. CI provides a reproducible measure of group-level effects on total ecosystem biomass (TEB), extending robustness evaluation beyond single-species removals.

### Definition

Let *B*^(0)^ denote the initial biomass vector of the intact food web. For a species set *K* = *{k*_1_*, . . ., k*_|_*_K_*_|_*}*, let TEB(*B*^(−^*^K^*^)^) be the equilibrium total ecosystem biomass after simultaneously removing all species in *K*, and let TEB(*B*^(−^*^kj^* ^)^) denote the equilibrium total ecosystem biomass after removing only species *k_j_*. The Cluster Influence of *K* is defined in Eq. (3).

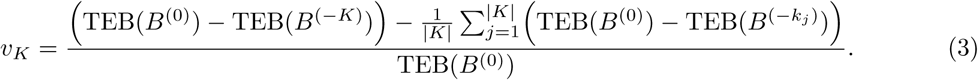

The first term reflects the observed impact of simultaneous removal of the group, while the second term represents the mean impact of removing its members individually. Averaging individual effects ensures comparability across species sets of different sizes. Thus, positive values of *v_K_* indicate stronger-than-mean group effects, consistent with synergistic influence, whereas negative values indicate weaker-than-mean group effects, consistent with redundancy or compensation. Values near zero indicate that the group effect is comparable to the mean individual effect.

### Sequential CI trajectories

To evaluate how cluster-level interactions accumulate during removal sequences, we compute CI for the prefixes of the greedy trajectories:

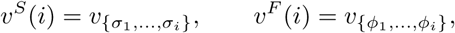

where (*σ*_1_*, . . ., σ_i_*) and (*ϕ*_1_*, . . ., ϕ_i_*) are the first *i* species removed along the SCT and FCT, respectively (Section 2.4.1). The resulting functions *v^S^*(*i*) and *v^F^* (*i*) constitute CI trajectories, which quantify how non-additive group effects change as removals proceed.

### Methodological considerations

CI is fully model-agnostic: it requires only the ability to simulate equilibrium biomass under simultaneous and individual removals. Because the computation for a set of size *i* requires *i* + 1 dynamic simulations, CI trajectories scale linearly with sequence length. In the context of MVSS identification, CI provides a complementary diagnostic by highlighting clusters in which joint removals induce disproportionately large functional losses. This enables a mechanistic link between minimal vital sets and measurable higher-order dependencies within the food web.

### 2.5 Structural and extinction-based comparison metrics

To compare DAR with existing robustness approaches, we also calculated LCC–AUC and SE metrics under several removal sequences. These comparison sequences were defined independently of biomass dynamics, except for the SCT and FCT. The degree-based sequence removed species in decreasing total degree. The in-degree-based sequence removed species in decreasing in-degree. The betweenness-based sequence removed species in decreasing betweenness centrality, and the PageRank-based sequence removed species in decreasing PageRank score. The random sequence removed species in a random order. We additionally evaluated the dynamically derived SCT and FCT so that topology-based and biomass-based removal sequences could be compared within the same AUC and SE calculations.

For the LCC–AUC analysis, the directed food web was converted to an undirected graph. Species were then removed sequentially according to each removal sequence, and after each removal step we recorded the size of LCC as a fraction of the original species richness. The AUC value was computed as the area under this LCC-retention curve using numerical trapezoidal integration. Lower AUC values indicate faster structural fragmentation, whereas higher AUC values indicate greater persistence of the largest connected component.

For the SE analysis, species removed according to each removal sequence were treated as primary removals. After each primary removal, species that no longer retained incoming trophic links in the remaining network were counted as secondary extinctions and removed before the next primary removal step. We recorded the cumulative fraction of primary removals and the cumulative fraction of secondary extinctions. The SE summary value was defined as the primary-removal fraction at which the combined fraction of primary and secondary losses first reached 50% of the original species richness. These metrics were used only as structural and extinction-based benchmarks for comparison with DAR.

### 2.6 Sensitivity and uncertainty analyses

To evaluate whether DAR estimates depended on uncertain bioenergetic parameters, we conducted parameter sensitivity and uncertainty analyses after computing the main DAR, MVSS, and CI metrics. We focused on four parameters: the Hill exponent (*h*), half-saturation density (*B*_0_), intraspecific interference strength (*i*^intra^), and predator–prey body-mass ratio (*Z*), which control functional-response nonlinearity, feeding saturation, consumer interference, and allometric metabolic scaling, respectively. Sensitivity was assessed using a one-at-a-time design, in which one parameter was varied while all others were held at their default values. Parameter uncertainty was then propagated using Latin Hypercube Sampling, with DAR, MVSS, and CI recalculated for each sampled parameter set.

## 3 Results

We first evaluated DAR, MVSS, and CI on virtual food webs generated under controlled gradients of species richness and connectance. We then compared DAR with structural fragmentation and secondary-extinction metrics to examine how biomass-based robustness relates to existing topology-based and extinction-based approaches. Finally, we assessed parameter sensitivity and uncertainty, before applying the framework to 16 empirical stream food webs as a demonstration on realistic food-web topologies.

### 3.1 Virtual food-web simulations

We first evaluated DAR on 120 virtual food webs generated under controlled gradients of species richness and connectance. The design included three richness levels (*S* = 20, 50, and 100), four connectance levels (*C* = 0.05, 0.10, 0.15, and 0.20), and 10 replicate webs for each *S × C* combination. DAR was computed for each web from the paired slowest-and fastest-collapse biomass trajectories. Figure 2 summarizes how biomass-based robustness varied across this controlled structural space; the grey band indicates the DAR range observed across the 16 empirical stream food webs.

**Figure 2:**
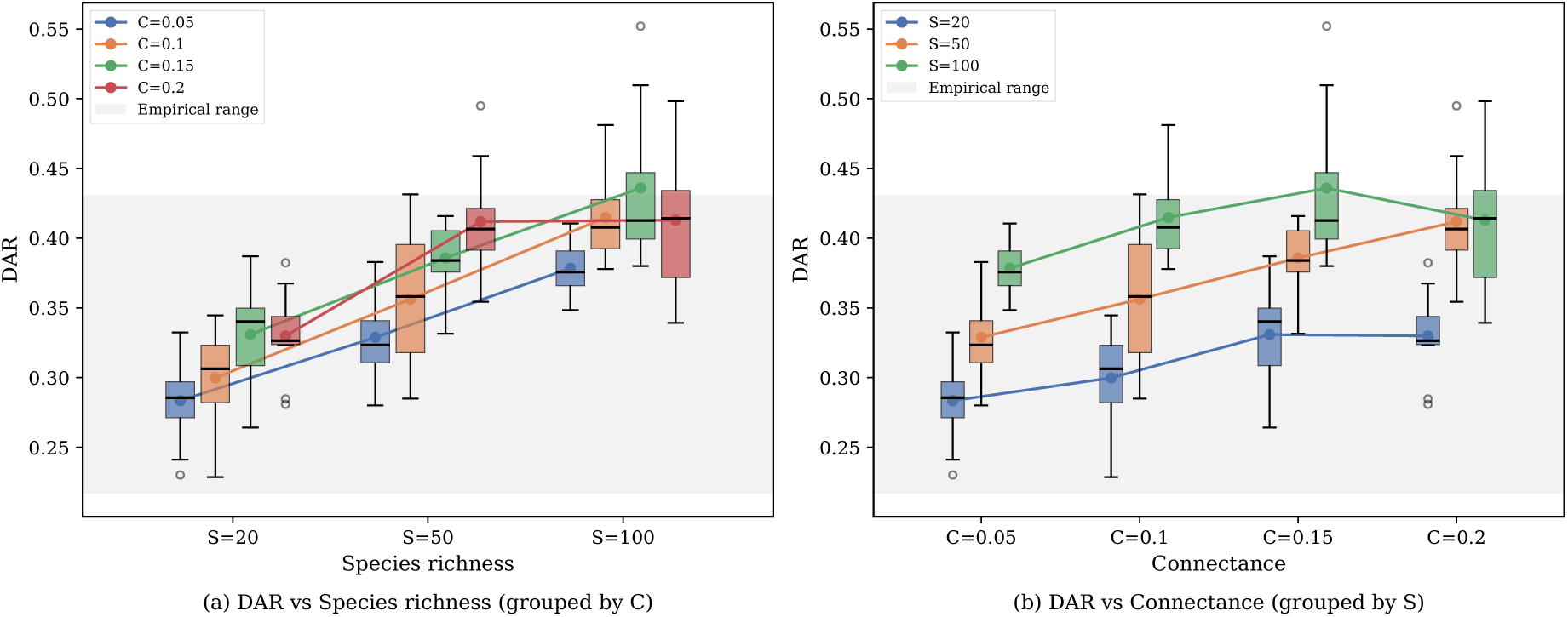
Dynamic Area-based Robustness (DAR) across virtual food webs generated under controlled gradients of species richness and connectance. (a) DAR grouped by species richness, with colours indicating connectance. (b) DAR grouped by connectance, with colours indicating species richness. Each box represents 10 replicate niche-model food webs for a given parameter combination, and lines connect group means. The grey band indicates the DAR range observed across the 16 empirical stream food webs.

DAR increased consistently with species richness. Across all four connectance levels, larger virtual food webs showed higher mean DAR than smaller webs, indicating that species-rich networks retained a broader biomass buffer before crossing the functional collapse threshold. Mean DAR increased from approximately 0.28–0.33 at *S* = 20, to 0.33–0.41 at *S* = 50, and to 0.38–0.44 at *S* = 100 depending on connectance.

The effect of connectance was positive but conditional on network size. At intermediate richness (*S* = 50), DAR increased monotonically with connectance, indicating that denser trophic interaction networks can retain more biomass under sequential species loss. However, this relationship was not universal. In the smallest webs (*S* = 20) and, most clearly, in the largest webs (*S* = 100), DAR increased from low to intermediate connectance but declined at the highest connectance level (*C* = 0.20), where mean DAR fell from 0.436 at *C* = 0.15 to 0.413 at *C* = 0.20. This non-monotonic response indicates that increasing interaction density does not always translate into greater dynamic robustness. Under such conditions, additional links may increase dependence on particular trophic configurations rather than simply adding functional redundancy.

Replicate variation further showed that DAR was not determined solely by the imposed values of *S* and *C*. Variation among replicate webs was modest in low-and intermediate-complexity settings, but became larger in species-rich, highly connected webs; the greatest replicate variability occurred in the *S* = 100, *C* = 0.15 and *C* = 0.20 treatments (SD *≈* 0.05). This indicates that, once structural complexity increases, the particular trophic arrangement generated by the niche model can strongly affect biomass-retention trajectories. Thus, DAR captures both broad structural effects, such as species richness and connectance, and network-specific dynamic responses arising from the realised topology.

The empirical food webs fell within the lower-to-intermediate portion of the virtual DAR range. This comparison suggests that the virtual simulations covered the empirical domain while also extending beyond it into higher-connectance and larger-network conditions. The virtual-web analysis therefore provides a controlled test of DAR across structural gradients broader than those available in the empirical dataset, while the empirical webs serve as realistic demonstrations of the same framework.

### 3.2 Comparison with structural and extinction-based robustness metrics

We next compared DAR with structural fragmentation and secondary-extinction metrics calculated on the same virtual food webs. The comparison included AUC and SE values under degree-, in-degree-, betweenness-, PageRank-, and random-removal sequences, as well as the biomass-derived SCT and FCT sequences, together with standard structural descriptors of the generated food webs.

DAR was strongly correlated with several existing robustness metrics (Fig. 3). The strongest associations were observed mainly with AUC-based targeted-removal metrics, especially Degree-AUC (*ρ* = 0.79) and Betweenness-AUC (*ρ* = 0.76). Among SE-based metrics, Betweenness-SE remained positively associated with DAR (*ρ* = 0.69), whereas Degree-SE was only moderately correlated (*ρ* = 0.60). These correlations indicate that, in controlled niche-model food webs, biomass-based robustness is most closely aligned with targeted-removal structural robustness, especially degree-and betweenness-based AUC. The association with secondary-extinction robustness is weaker and depends strongly on the removal sequence. However, the associations were not uniform across all metrics. DAR was weakly correlated with PageRank-SE (*ρ* = 0.29), SCT-SE (*ρ* = 0.10), and Random-SE (*ρ* = 0.40), and was negatively correlated with FCT-SE (*ρ* = *−*0.31). This indicates that the relationship between biomass-based robustness and binary secondary-extinction robustness is substantially weaker than previously suggested and depends strongly on the removal sequence.

**Figure 3:**
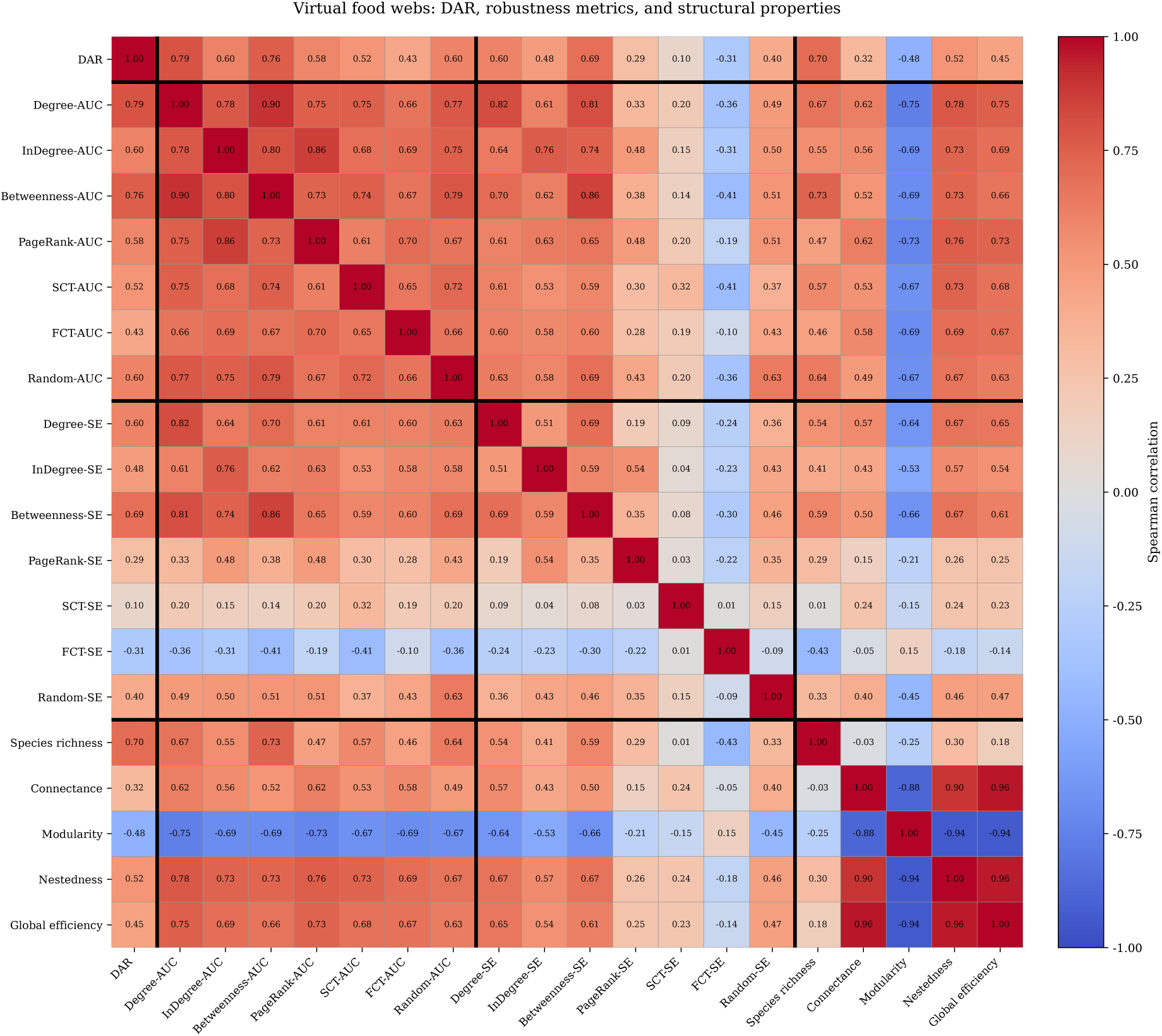
Spearman correlations among DAR, AUC-based robustness, SE-based robustness, and structural descriptors across 120 virtual food webs. Black lines separate DAR, AUC metrics, SE metrics, and structural properties.

Notably, AUC values derived from the biomass-based SCT and FCT sequences were moderately correlated with DAR (SCT-AUC *ρ* = 0.52, FCT-AUC *ρ* = 0.43), whereas the corresponding SE values showed little or opposite association with DAR (SCT-SE *ρ* = 0.10, FCT-SE *ρ* = *−*0.31). Thus, even when the removal sequence is biomass-derived, binary secondary-extinction summaries do not reproduce biomass-retention robustness.

DAR was also strongly associated with species richness (*ρ* = 0.70) and moderately associated with nestedness (*ρ* = 0.52) and global efficiency (*ρ* = 0.45), while showing only a weak association with connectance (*ρ* = 0.32) and a negative association with modularity (*ρ* = *−*0.48). These structural descriptors were themselves highly correlated, particularly connectance, nestedness, and global efficiency. Therefore, the observed relationships should not be interpreted as independent effects of individual structural properties. Rather, they show that DAR covaries with broader gradients of food-web complexity and integration in the virtual networks.

DAR recovered part of the variation captured by AUC-based robustness metrics and by some targeted-removal SE metrics, especially Betweenness-SE. However, the correlation pattern varied strongly across metric classes. The strongest correlations occurred for Degree-AUC and Betweenness-AUC, whereas weaker or even negative associations were observed for PageRank-SE, SCT-SE, FCT-SE, and Random-SE. Thus, DAR is related to established structural robustness measures, but it is not reducible to either AUC-based fragmentation or binary secondary-extinction summaries.

### 3.3 MVSS and CI across virtual food webs

Across the 120 niche-model food webs, MVSS size and MVSS proportion varied systematically with network structure (Figure 4A). The mean MVSS proportion across all *S × C* groups was 0.298, with group means ranging from 0.154 to 0.501, and mean MVSS size ranging from 5.3 to 50.1 species. For *S* = 20 and *S* = 50, MVSS proportion generally declined with connectance, indicating that more connected virtual webs tended to maintain function with a smaller retained core. The *S* = 100, *C* = 0.20 treatment showed the largest variance, indicating that high-richness, high-connectance replicate webs can deviate substantially from this monotonic trend.

**Figure 4:**
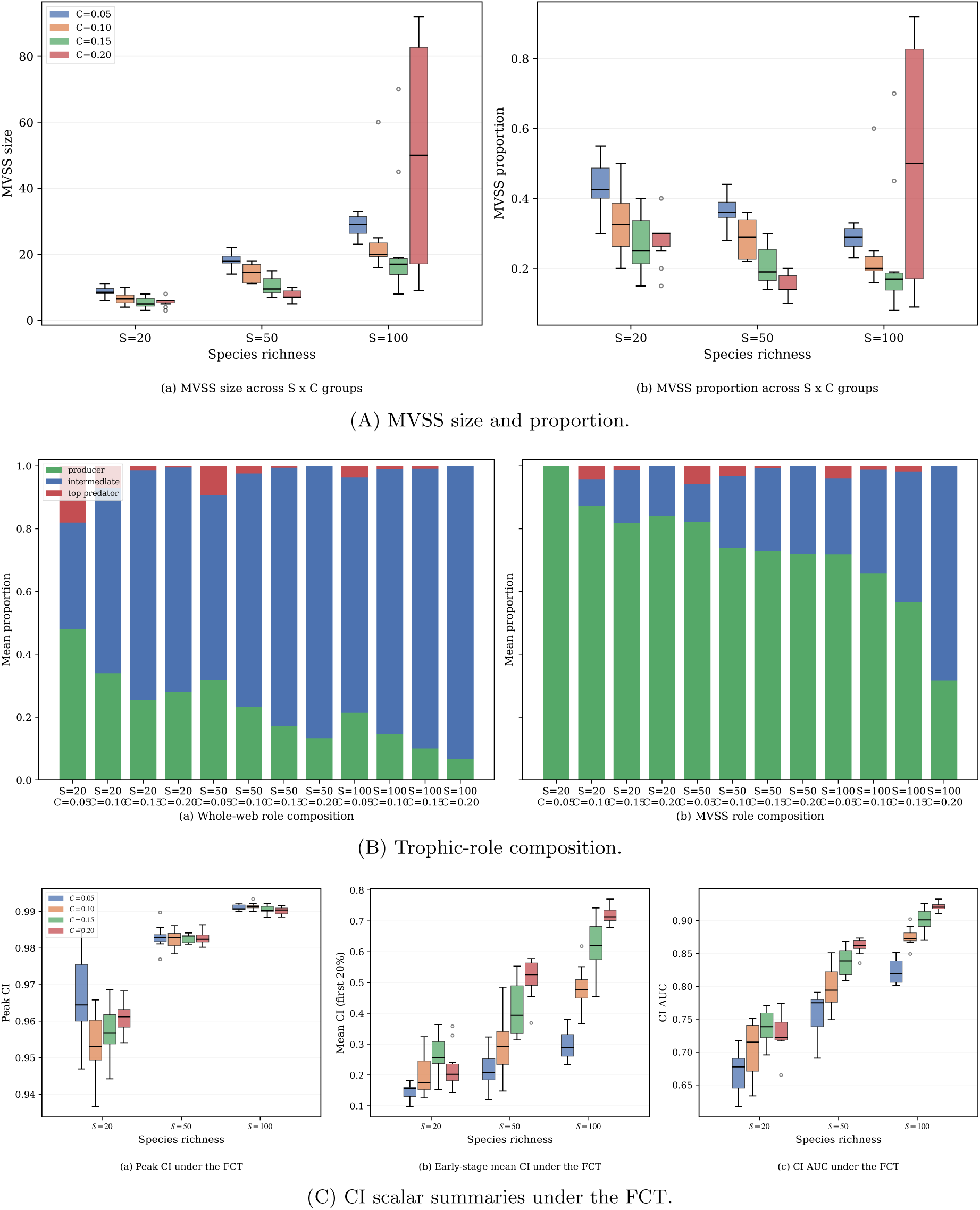
MVSS and CI across virtual food webs. Panel (A) summarises MVSS size and MVSS proportion across the 120 niche-model food webs grouped by species richness and connectance. Panel (B) compares the mean trophic-role composition of whole virtual food webs and their MVSS subsets. Panel (C) summarises the distributions of key CI statistics under the FCT. Additional broad trophic-class, grouped-trajectory, and MVSS–CI linkage results are shown in Figures S2– S4.

The trophic composition of the MVSS differed strongly from that of the whole webs (Figure 4B). In the full virtual webs, intermediate consumers dominated (73.4% on average), whereas producers accounted for 22.8% and top predators for only 3.8%. In the MVSS, this pattern reversed: producers increased to 73.3%, intermediates declined to 24.8%, and top predators to 1.9%. Under the same prey-averaged trophic-level definition used elsewhere in this study (following Levine (1980) and Williams and Martinez (2004)), the broad trophic-class summary led to the same conclusion. Whole webs contained on average 22.8% basal producers, 14.6% low-level consumers, 24.1% mid-level consumers, and 38.4% higher-level consumers, whereas MVSS subsets contained 73.3% basal producers, 0.6% low-level consumers, 5.7% mid-level consumers, and 20.3% higher-level consumers (Figure S2). Thus, the minimal sufficient sets in the virtual webs were strongly enriched in basal species, with a smaller retained contribution from higher trophic positions.

Virtual webs also showed a clear contrast in group-level interaction effects. Under the SCT, early-stage CI was typically negative, with an across-group mean of *−*0.169, indicating weaker-than-mean group effects or compensation among the species removed early in the least-damaging trajectory. Under the FCT, early-stage CI was consistently positive, with a corresponding mean of 0.367, indicating stronger-than-mean group effects among the species removed early in the fast-collapse trajectory. This contrast remained evident in the scalar summaries (Figure 4C): the mean CI area under the curve was 0.801 under the FCT but *−*0.136 under the SCT.

Moreover, webs with smaller MVSS fractions exhibited stronger CI. MVSS proportion was negatively associated with peak CI (*ρ* = *−*0.183, *p* = 4.56 *×* 10^−2^), early-stage CI (*ρ* = *−*0.529, *p* = 5.40 *×* 10^−10^), CI area (*ρ* = *−*0.480, *p* = 2.91 *×* 10^−8^), and, most strongly, CI at the critical collapse step (*ρ* = *−*0.614, *p* = 9.00 *×* 10^−14^). The grouped CI trajectories and the MVSS–CI linkage plots are shown in Figures S3–S4.

Taken together, these results show that smaller MVSS fractions were associated with stronger positive CI under the FCT, whereas the SCT was generally associated with negative or weaker group-level effects during early removals.

### 3.4 Sensitivity and uncertainty analyses

We next assessed whether the core DAR results were robust to uncertainty in the bioenergetic parameterization. Across 12 representative virtual food webs spanning all *S × C* combinations, the strongest one-at-a-time sensitivity effects arose from the predator–prey body-mass ratio and the half-saturation density, followed by the Hill exponent, whereas intraspecific interference had a weaker influence overall. Averaged over the representative networks, varying each parameter changed DAR by between *−*7.1% and +30.6% for *Z*, between *−*10.0% and +28.4% for *B*_0_, between *−*11.4% and +19.8% for *h*, and between *−*2.4% and +2.5% for *i*^intra^ (Figure 5a).

**Figure 5:**
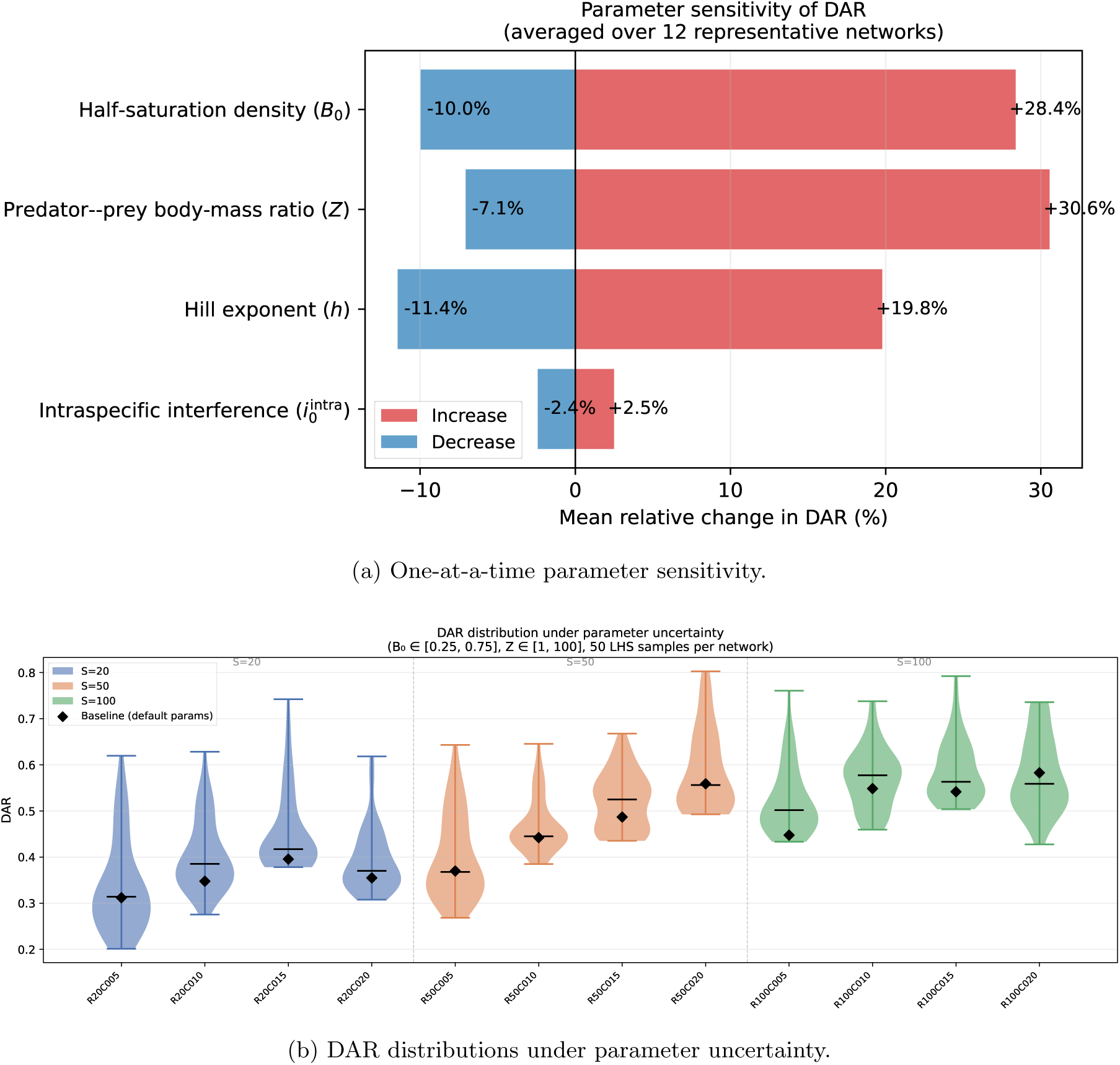
Sensitivity and uncertainty analyses for DAR. Panel (a) shows the mean relative changes induced by one-at-a-time perturbations of the four focal bioenergetic parameters. Panel (b) shows the distribution of DAR values obtained under Latin Hypercube parameter sampling. Additional sensitivity and uncertainty plots are provided in Figure S5 and Figure S6.

To propagate uncertainty more formally, we sampled *B*_0_ and *Z* using Latin Hypercube Sampling and recomputed DAR for each sampled parameter set. The resulting DAR distributions remained structured by network size and connectance, while showing substantial but interpretable spread (Figure 5b). Across the 12 representative structural treatments, the interquartile range of DAR varied from 0.046 to 0.134. Thus, the exact numerical value of DAR depends on bioenergetic assumptions, but the main structural contrasts identified in the virtual-web analysis remain robust. Additional sensitivity curves and uncertainty-width plots are provided in Figures S5–S6.

### 3.5 Empirical food-web demonstration

We finally applied the framework to 16 empirical stream food webs as a demonstration on realistic interaction topologies. The empirical results were qualitatively consistent with the virtual-food-web analysis, but quantitatively narrower and ecologically more heterogeneous. Empirical DAR values ranged from 0.217 to 0.431, placing the stream webs within a restricted subset of the broader virtual range. The empirical DAR values therefore fell within the range covered by the virtual simulations, while the virtual webs also extended into larger and more highly connected structural regimes not represented in the empirical dataset.

Across the 16 stream webs, the mean MVSS proportion was 0.582, ranging from 0.327 to 0.785, which was larger than the average proportion observed in the virtual food webs (0.298). Thus, the empirical stream webs generally required a larger retained species fraction to remain above the collapse threshold than the virtual niche-model webs.

Likewise, the empirical CI trajectories preserved the same directional contrast observed in the virtual webs: early-stage CI was negative under the SCT (*−*0.238) but positive under the FCT (0.131), and the cumulative CI area was much larger under the FCT (0.651 versus 0.006). The empirical analyses reproduced the main directional CI pattern observed in the virtual webs, while showing larger MVSS proportions and more system-specific variation. The empirical MVSS trophic-class composition is shown in Figure S9, and additional empirical AUC and SE trajectories are provided in Figures S7 and Figure S8.

## 4 Discussion

We developed a biomass-dynamic framework for evaluating food-web robustness under progressive species loss. The framework combines Dynamic Area-based Robustness (DAR), Minimal Vital Species Set (MVSS), and Cluster Influence (CI) to connect three levels of analysis: system-level biomass retention, collapse-associated species sets, and group-level effects of species removals. The virtual-food-web simulations showed that DAR increased consistently with species richness, which acted as a more uniform predictor of dynamic robustness than connectance. The effect of connectance was instead non-monotonic in the most species-rich webs, where the highest connectance level reduced rather than increased DAR; this suggests that dense interaction networks can concentrate dependence on particular trophic pathways rather than simply adding functional redundancy. The empirical stream food webs fell within a narrower, lower-to-intermediate portion of the virtual range. Together, these results indicate that the framework can be used both for controlled structural comparisons and for demonstration on realistic food-web topologies.

The comparison with AUC-and SE-based metrics clarifies what DAR adds to existing robustness analyses (Albert et al., 2000; Schneider et al., 2011; Dunne et al., 2004; Dunne and Williams, 2009). DAR was correlated with several structural robustness metrics, particularly Degree-AUC and Betweenness-AUC, showing that biomass-based robustness is not detached from structural vulnerability. However, the correlations were not uniform across removal sequences or metric classes. The strongest associations were found for AUC-based metrics under degree-and betweenness-based removals, whereas the associations with SE-based metrics were weaker and less consistent. Although Degree-SE and Betweenness-SE were still positively correlated with DAR, PageRank-SE, SCT-SE, and Random-SE showed weaker associations, and FCT-SE was negatively correlated with DAR. This indicates that DAR is not simply a rescaled version of structural fragmentation or SE-based robustness. AUC primarily summarizes how rapidly the network fragments under species removal, whereas SE evaluates whether consumers lose trophic resources and become secondarily extinct. DAR instead tracks continuous biomass-retention trajectories relative to an explicit functional collapse threshold. Notably, the AUC values computed along the biomass-derived SCT and FCT removal sequences were only moderately correlated with DAR, whereas the corresponding SE values were weakly or negatively correlated with DAR. This shows that structural fragmentation, secondary-extinction responses, and biomass retention capture distinct aspects of robustness even when the removal sequence is biomass-derived. Therefore, DAR complements, rather than replaces, existing structural and SE-based robustness metrics. It is related to established AUC-based robustness measures, especially under degree-and betweenness-based removals, but it captures biomass-retention patterns that are not recovered by either structural fragmentation summaries or SE-based indices.

The MVSS and CI results further show that robustness can depend on species groups rather than only on individually central species. In the virtual food webs, MVSS subsets were strongly enriched in basal species, indicating that persistence above the biomass-collapse threshold depended heavily on maintaining basal energy input under the present model parameterization. This result is consistent with bottom-up trophic control, where primary producers and basal resources constrain the biomass that can be supported at higher trophic levels (Pace et al., 1999; Hooper et al., 2005). At the same time, the empirical stream food webs required larger retained species fractions, suggesting that realistic trophic heterogeneity, specialization, and system-specific interaction structure can reduce the extent to which a small basal-dominated set is sufficient. CI provides an additional interpretation of these retained sets by quantifying whether the simultaneous removal of a species group produces a stronger or weaker effect than the mean effect of removing its members individually. Negative CI under the SCT is consistent with functional redundancy or compensation, where species removed early have overlapping or weakly unique effects on TEB. Positive CI under the FCT indicates stronger-than-mean group effects (Loreau and Hector, 2001; Ball et al., 2008; Guo et al., 2025), suggesting that collapse-prone species sets can act as tightly coupled functional groups. Moreover, webs with smaller minimal vital sets exhibited stronger positive CI, directly linking the two metrics: a more compact collapse-associated set tended to act as a more tightly coupled, synergistic functional group rather than a collection of independently acting species. In this sense, the framework extends single-species notions of keystone effects toward the identification of collapse-associated species sets and their group-level influence.

Several caveats should be considered when interpreting the proposed framework. First, DAR is a parameterized measure of biomass-based functional resistance, not a universal measure of ecosystem stability. It depends on the chosen bioenergetic model, parameter values, initial biomass conditions, and the definition of the collapse threshold. The sensitivity and uncertainty analyses showed that DAR was especially influenced by the predator–prey body-mass ratio and half-saturation density, even though the main structural contrasts remained evident. Second, MVSS is identified from a heuristic fastest-collapse trajectory rather than from an exhaustive search over all possible species subsets. It should therefore be interpreted as an algorithmically identified collapse-associated vital set, not as a guaranteed globally minimal set. Third, CI measures group-level effects on TEB, but it does not by itself identify the underlying ecological mechanism. Positive or negative CI may arise from trophic complementarity, redundancy, indirect effects, compensation, or the particular parameterization of consumer–resource interactions. Fourth, the empirical analyses were intended as a demonstration on realistic stream food-web topologies, whereas the primary inference came from virtual food webs generated under controlled structural gradients. Broader empirical validation across additional ecosystems, weighted interactions, and independently estimated biomass data will be needed before drawing general ecological conclusions.

Despite these limitations, the DAR–MVSS–CI framework provides a reproducible way to connect food-web structure with biomass-based functional collapse. By distinguishing structural disassembly, binary secondary extinctions, and dynamic biomass retention, the framework helps clarify when topology-based robustness is informative and when dynamic simulations provide additional diagnostic resolution. Future work could extend the framework by incorporating interaction strengths, adaptive rewiring, environmental forcing, trait-dependent parameters, or alternative ecosystem functions beyond total biomass. These extensions would allow the same general approach to evaluate functional resistance under a wider range of disturbance scenarios and food-web models.

