## Supporting Information for "Functional Robustness of Food Webs: A Dynamic Biomass Framework with Vital Species Sets and Cluster Influence"

Additional details of the article are provided here.

#### Supplementary for model

##### Supplementary for parameter $t$ sensitivity

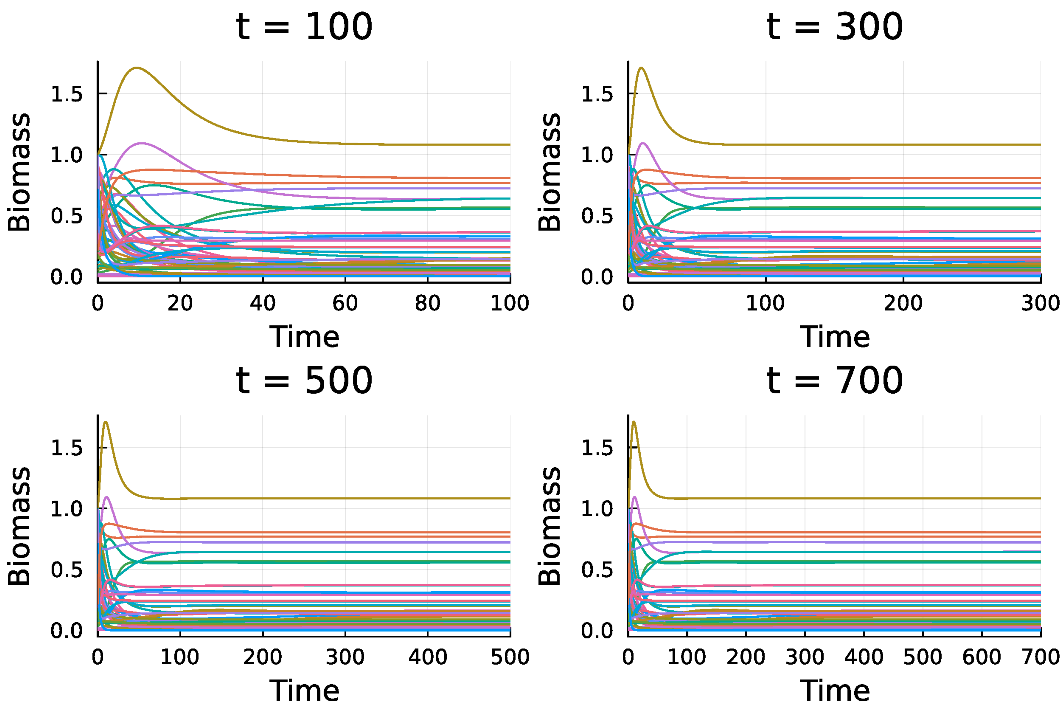


Figure S1: Sensitivity of biomass trajectories to simulation horizon $t$. Each panel shows the temporal dynamics of a random 50-species niche-model food web simulated up to different time horizons. Results indicate that horizons of $t \geq 500$ are sufficient for convergence.

##### Supplementary for model parameters

Table S1: Default parameters for the bioenergetic model.

| **Parameters** | **Symbol** | **Default producers** | **Default consumers** | **References** |
|---|---|---|---|---|
| Body mass | $M_i$ | 1 | $Z^{TL_i - 1}$ | (Brose et al., 2006) |
| Intrinsic growth | $r_i$ | $M_i^{-0.25}$ | 0 | (Miele et al., 2019) |
| Carrying capacity | $K_i$ | $M_i^{0.25}$ | $\emptyset$ | (Miele et al., 2019) |
| Metabolism | $x_i$ | 0 | $\begin{cases} 0.314 M_i^{-0.25}, & \text{if invertebrates} \\ 0.88 M_i^{-0.25}, & \text{if vertebrates} \end{cases}$ | (Delmas et al., 2017; Miele et al., 2019) |
| Maximum consumption | $y_i$ | 0 | $\begin{cases} 8, & \text{if invertebrates} \\ 4, & \text{if vertebrates} \end{cases}$ | (Delmas et al., 2017) |
| Mortality | $d_i$ | 0 | $0.0314 M_i^{-0.25}$ | (Miele et al., 2019) |
| Preference | $\omega_{ij}$ | $\emptyset$ | $\frac{1}{\lvert\{\text{prey}_i\}\rvert}$ | (Delmas et al., 2017; Miele et al., 2019) |
| Efficiency | $e_{ij}$ | $\emptyset$ | $\begin{cases} 0.45, & \text{herbivores} \\ 0.85, & \text{carnivores} \end{cases}$ | (Delmas et al., 2017; Miele et al., 2019) |
| Half-saturation density | $B_0$ | $\emptyset$ | 0.5 | (Delmas et al., 2017) |
| Hill exponent | $h$ | $\emptyset$ | 2 | (Delmas et al., 2017) |
| Intraspecific interference | $i_0^{\text{intra}}$ | $\emptyset$ | 0 | (Delmas et al., 2017) |

### Supplementary for virtual food webs

#### Virtual food webs broad trophic composition

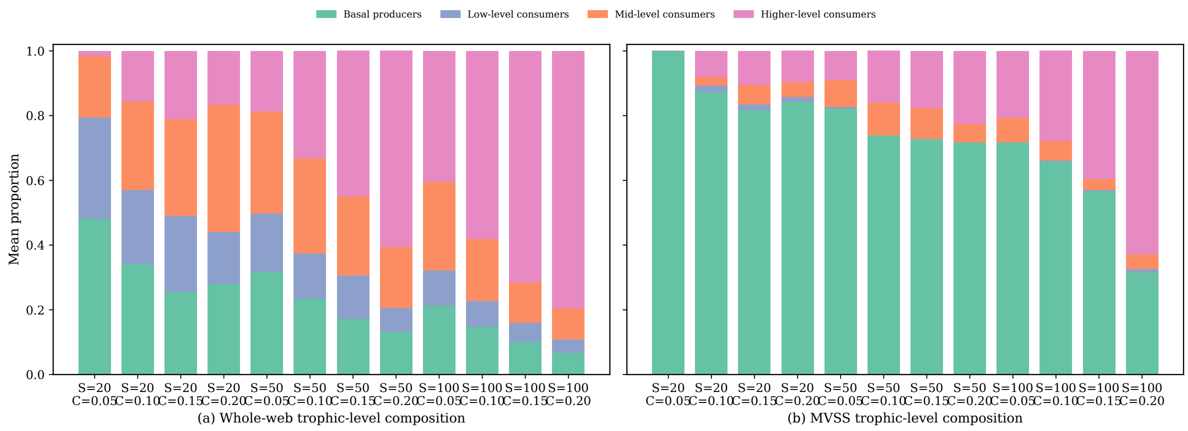


Figure S2: Broad trophic-class composition of virtual food webs and their MVSS subsets. Species were assigned prey-averaged trophic levels (following Levine (1980) and Williams and Martinez (2004)) and then grouped into broad trophic classes for display. The left panel shows the mean composition of the full virtual food webs, and the right panel shows the corresponding composition of the MVSS subsets, both summarized within each $S \times C$ group.

#### Virtual food webs CI trajectories

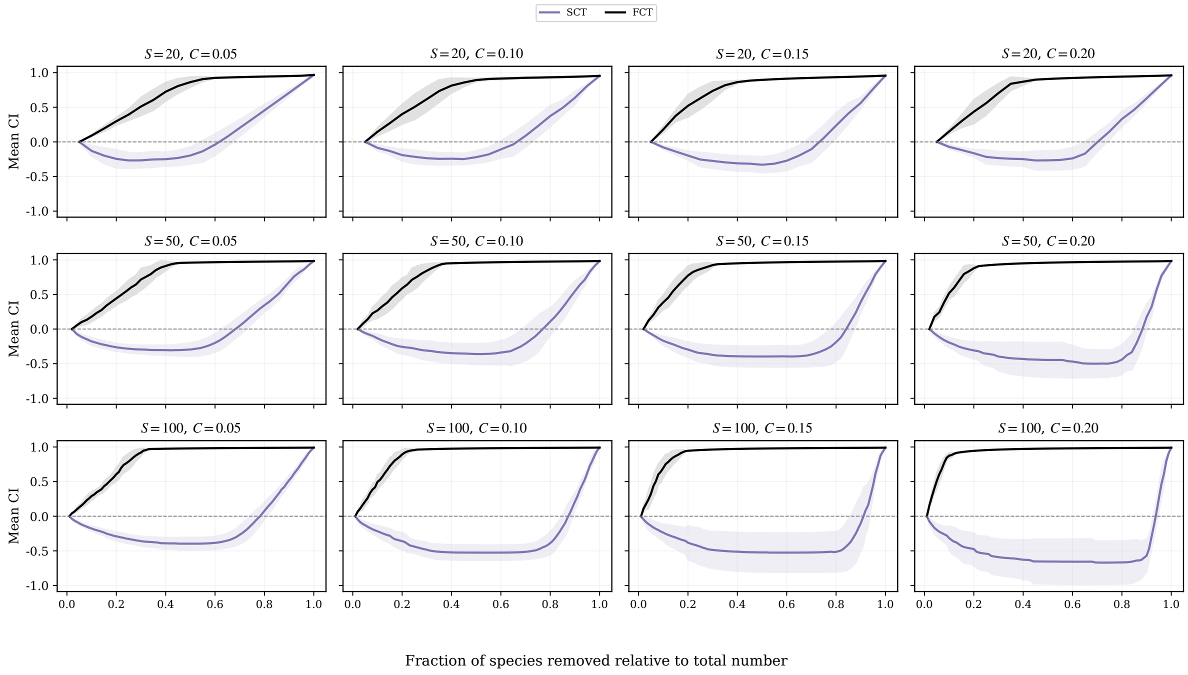


Figure S3: Grouped CI trajectories across the 120 virtual food webs. Each panel corresponds to one $S \times C$ treatment and shows the mean trajectory across 10 replicate webs, with shaded bands indicating one standard deviation. The purple line represents the slowest-collapse trajectory (SCT), and the black line represents the fastest-collapse trajectory (FCT).

#### Virtual food webs of MVSS and CI

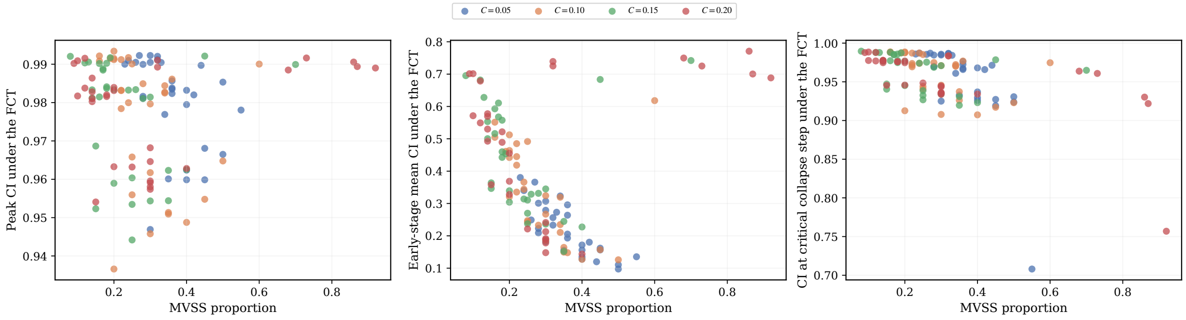


(a) MVSS proportion vs peak CI
Spearman $\rho$ = -0.18

(b) MVSS proportion vs early-stage CI
Spearman $\rho$ = -0.53

(c) MVSS proportion vs critical-step CI
Spearman $\rho$ = -0.61

Figure S4: Relationships between MVSS proportion and FCT-based CI strength across virtual food webs. Each point represents one virtual web. The panels compare MVSS proportion with peak CI, early-stage mean CI, and CI at the critical collapse step under the FCT.

#### Virtual food webs sensitivity analysis

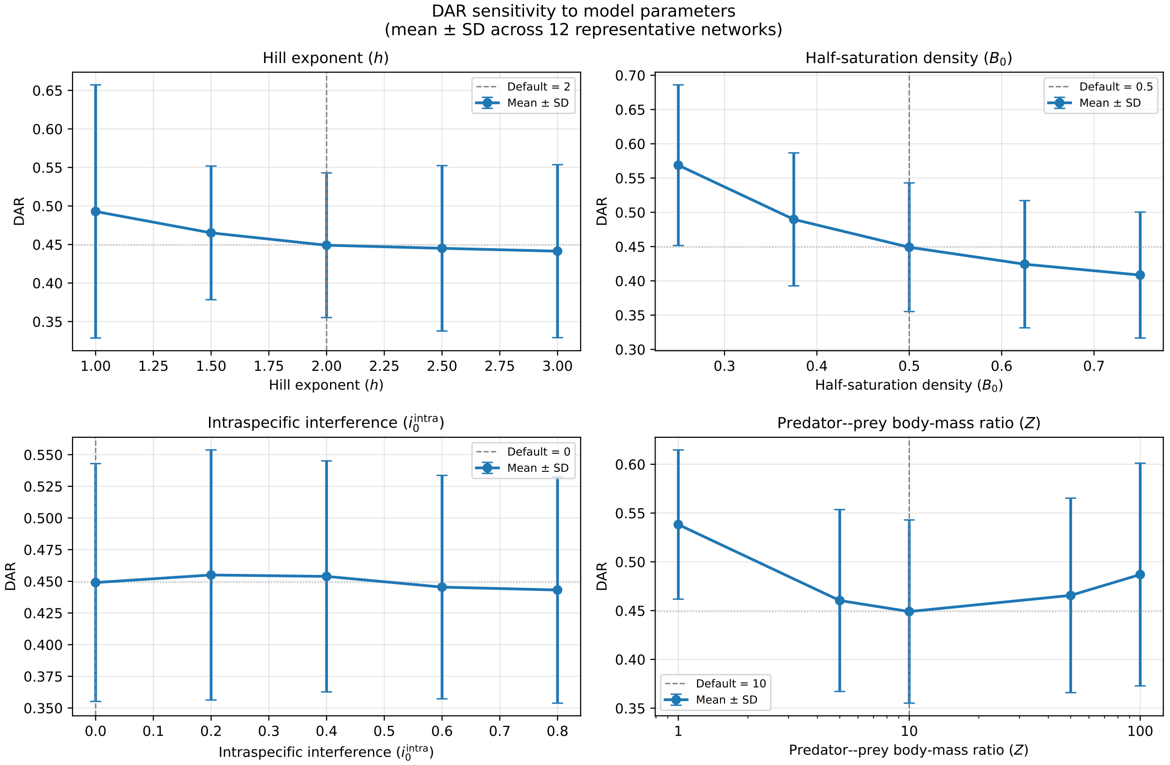


Figure S5: Detailed sensitivity curves for DAR across representative virtual food webs. Each panel summarizes how DAR changes as one focal bioenergetic parameter is varied while the others are held at their default values. Error bars denote the across-network spread for the 12 representative virtual food webs.

#### Virtual food webs uncertainty analysis

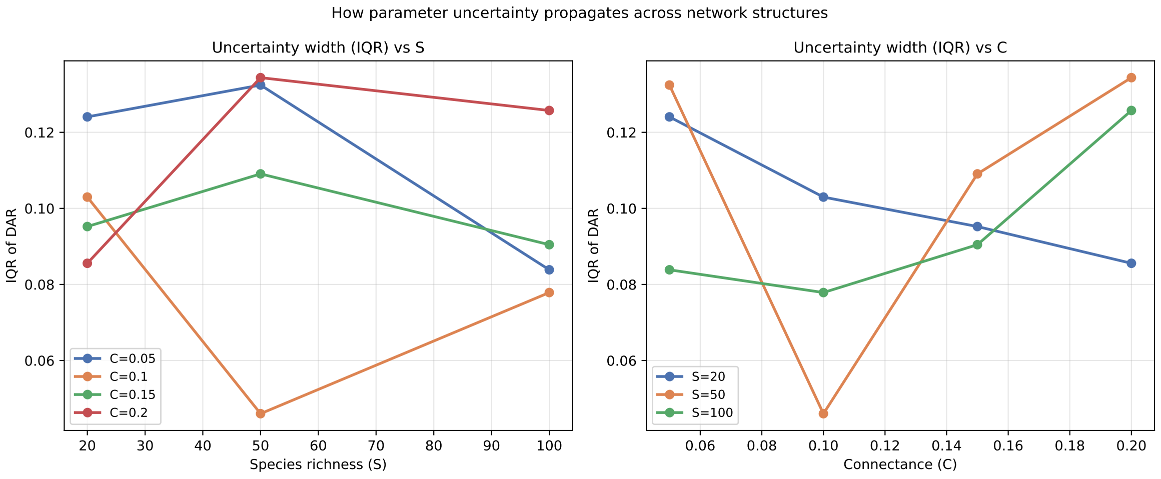


Figure S6: Propagation of parameter uncertainty across representative virtual food webs. The left panel shows how the interquartile range of DAR changes with species richness ($S$) when grouped by connectance ($C$), and the right panel shows the corresponding connectance dependence when grouped by species richness.

### Supplementary for empirical food webs

#### Empirical food webs structural properties

Modularity was computed after symmetrizing the directed food web and applying the Louvain algorithm over 1000 independent runs with different random seeds; the partition with the highest modularity was retained for reporting. Global efficiency was calculated as the mean inverse shortest-path length, with disconnected pairs assigned zero contribution.

Table S2: Structural properties and surrounding catchment context of the 16 empirical stream food webs. The food webs are aquatic stream food webs; the catchment labels describe the surrounding vegetation or land-use context rather than terrestrial food-web habitats. $S$ denotes species richness, $C$ connectance, $M$ modularity, and $GE$ global efficiency.

| **Food web** | **Surrounding Catchment** | **S**† | **C**‡ | **M**§ | **GE**¶ | **References** |
|---|---|---|---|---|---|---|
| AkatoreB | Pine forest | 58 | 0.0348 | 0.3320 | 0.3831 | (Thompson and Townsend, 2005) |
| Blackrock | Pasture grassland | 87 | 0.0495 | 0.2305 | 0.4581 | (Thompson and Townsend, 2005) |
| Broad | Pasture grassland | 95 | 0.0626 | 0.1813 | 0.4704 | (Thompson and Townsend, 2005) |
| Catlins | Pine forest | 49 | 0.0458 | 0.2744 | 0.4511 | (Thompson and Townsend, 2005) |
| DempstersAu | Tussock grassland | 86 | 0.0561 | 0.2272 | 0.4450 | (Thompson and Townsend, 1999) |
| DempstersSp | Tussock grassland | 97 | 0.0572 | 0.2016 | 0.4387 | (Thompson and Townsend, 1999) |
| German | Tussock grassland | 86 | 0.0477 | 0.2794 | 0.4300 | (Thompson et al., 2001) |
| Kyeburn | Tussock grassland | 98 | 0.0655 | 0.1811 | 0.4919 | (Thompson et al., 2001) |
| LilKyeburn | Tussock grassland | 78 | 0.0616 | 0.2634 | 0.4712 | (Thompson et al., 2001) |
| Martins | Pine forest | 105 | 0.0311 | 0.3339 | 0.4281 | (Thompson and Townsend, 2003) |
| NorthCol | Broadleaf forest | 78 | 0.0396 | 0.3186 | 0.4556 | (Thompson and Townsend, 2005) |
| Powder | Broadleaf forest | 78 | 0.0440 | 0.2694 | 0.4508 | (Thompson and Townsend, 2005) |
| Stony | Tussock grassland | 113 | 0.0652 | 0.1618 | 0.4860 | (Thompson et al., 2001) |
| SuttonSp | Tussock grassland | 79 | 0.0627 | 0.1698 | 0.4347 | (Thompson and Townsend, 1999) |
| Troy | Pine forest | 78 | 0.0298 | 0.3651 | 0.3923 | (Thompson and Townsend, 2003) |
| Venlaw | Pine forest | 69 | 0.0393 | 0.2923 | 0.4069 | (Thompson and Townsend, 2003) |

#### Empirical food webs AUC results

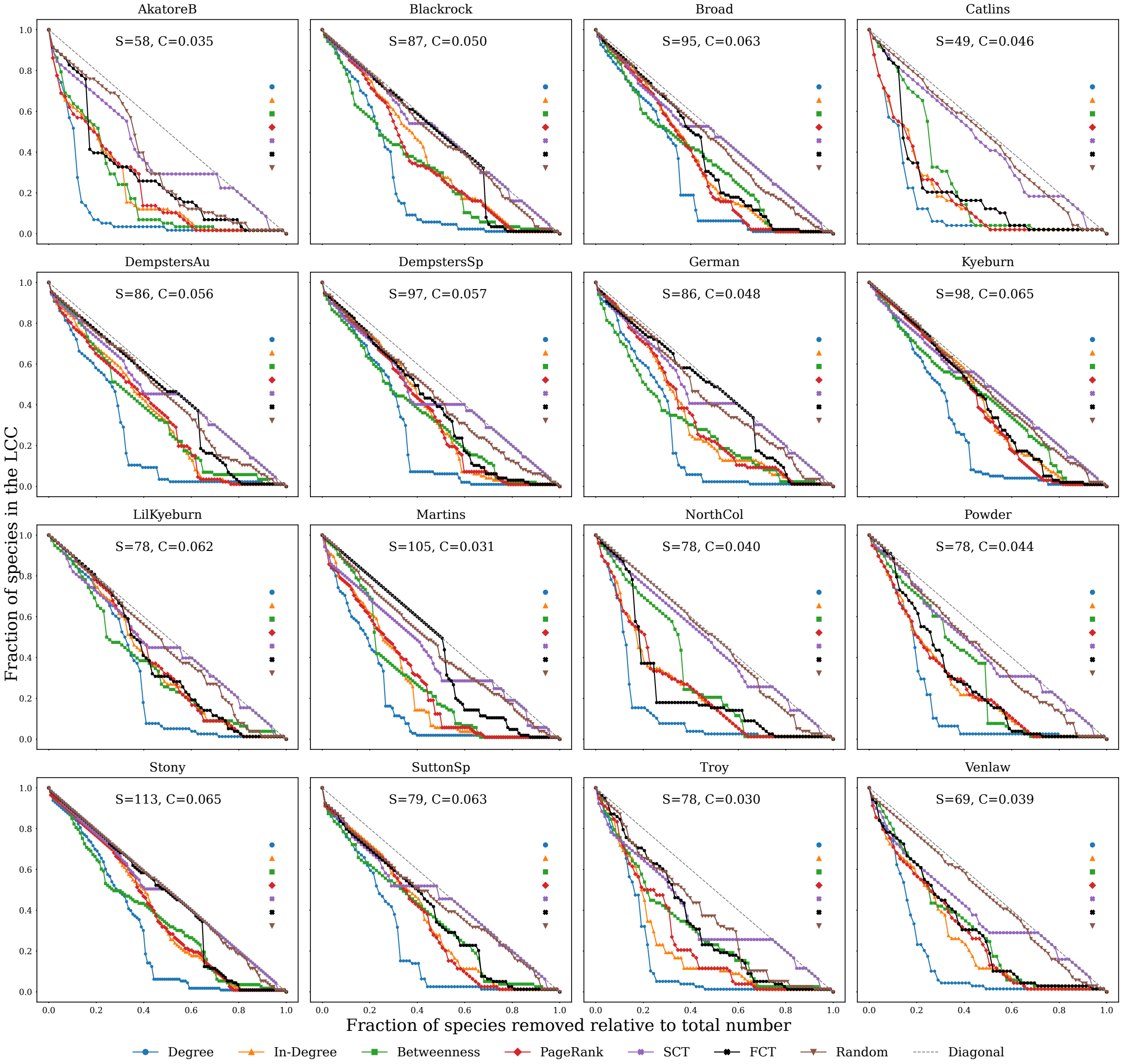


Figure S7: LCC size trajectories in the 16 empirical stream food webs under structural, dynamic, and random removal sequences. Each subplot represents one empirical stream food web. The x-axis denotes the fraction of species removed relative to the total number of species, and the y-axis represents the fraction of species retained in the largest connected component (LCC). Species richness ($S$) and connectance ($C$) are annotated in each subplot. The diagonal line is shown as a reference.

Table S3: AUC values for the 16 empirical stream food webs under structural, dynamic, and random removal sequences. Columns $D_{\mathrm{AUC}}$, $ID_{\mathrm{AUC}}$, $BET_{\mathrm{AUC}}$, $PR_{\mathrm{AUC}}$, $SCT_{\mathrm{AUC}}$, $FCT_{\mathrm{AUC}}$, and $\mathrm{Random}_{\mathrm{AUC}}$ denote degree-based, in-degree-based, betweenness-based, PageRank-based, SCT-based, FCT-based, and random removal sequences, respectively. Bold values indicate the top three values within each AUC metric. Larger values indicate higher robustness. The average row represents the average structural disassembly effect of each sequence.

| **Food web** | **D_AUC** | **ID_AUC** | **BET_AUC** | **PR_AUC** | **SCT_AUC** | **FCT_AUC** | **Random_AUC** |
|---|---|---|---|---|---|---|---|
| AkatoreB | 0.1130 | 0.2173 | 0.2084 | 0.2265 | 0.3977 | 0.2818 | 0.4090 |
| Blackrock | 0.2404 | **0.3814** | 0.3149 | 0.3601 | **0.4816** | **0.4483** | 0.4342 |
| Broad | 0.2736 | 0.3546 | **0.3565** | 0.3382 | 0.4707 | 0.3859 | 0.4676 |
| Catlins | 0.1372 | 0.1855 | 0.2584 | 0.1880 | 0.4429 | 0.2214 | 0.4663 |
| DempstersAu | 0.2339 | 0.3451 | 0.3488 | 0.3434 | 0.4594 | 0.4373 | **0.4698** |
| DempstersSp | 0.2615 | 0.3609 | 0.3374 | 0.3546 | 0.4417 | 0.3844 | 0.4457 |
| German | 0.2370 | 0.3342 | 0.2919 | 0.3407 | 0.4498 | **0.4484** | 0.4593 |
| Kyeburn | **0.2815** | **0.4185** | **0.4210** | **0.3966** | **0.4768** | 0.4166 | **0.4717** |
| LilKyeburn | **0.3008** | 0.3795 | 0.3506 | **0.3884** | 0.4581 | 0.3918 | 0.4528 |
| Martins | 0.1859 | 0.2610 | 0.2881 | 0.2769 | 0.4151 | 0.4283 | 0.4384 |
| NorthCol | 0.1435 | 0.2410 | 0.3203 | 0.2446 | 0.4675 | 0.2554 | 0.4666 |
| Powder | 0.1885 | 0.2673 | 0.3319 | 0.2704 | 0.4451 | 0.2926 | 0.4599 |
| Stony | **0.2862** | **0.3864** | **0.3635** | **0.3872** | **0.4821** | **0.4466** | **0.4677** |
| SuttonSp | 0.2356 | 0.3487 | 0.3515 | 0.3372 | 0.4494 | 0.3771 | 0.4349 |
| Troy | 0.1650 | 0.2265 | 0.2972 | 0.2536 | 0.3782 | 0.3363 | 0.4236 |
| Venlaw | 0.1629 | 0.2587 | 0.3030 | 0.2780 | 0.3965 | 0.3013 | 0.4187 |
| **Average** | 0.2154 | 0.3104 | 0.3215 | 0.3115 | 0.4445 | 0.3658 | 0.4491 |

#### Empirical food webs SE results

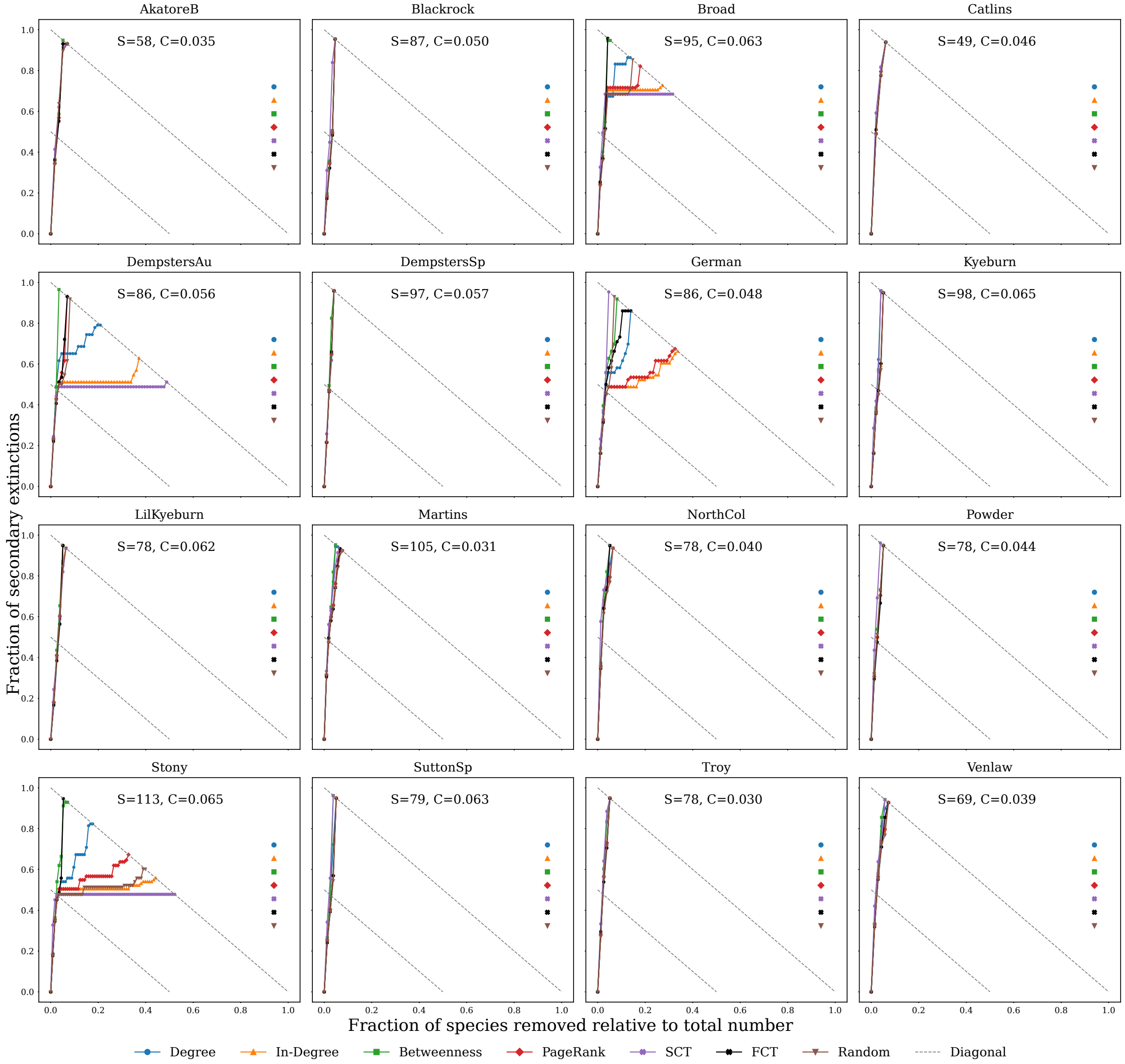


Figure S8: Secondary-extinction trajectories in the 16 empirical stream food webs under structural, dynamic, and random removal sequences. Each subplot represents one empirical stream food web. The x-axis denotes the fraction of species removed relative to the total number of species, and the y-axis represents the fraction of secondary extinctions. Species richness ($S$) and connectance ($C$) are annotated in each subplot. The diagonal line is shown as a reference.

Table S4: SE values for the 16 empirical stream food webs under structural, dynamic, and random removal sequences. Columns $D_{\mathrm{SE}}$, $ID_{\mathrm{SE}}$, $BET_{\mathrm{SE}}$, $PR_{\mathrm{SE}}$, $SCT_{\mathrm{SE}}$, $FCT_{\mathrm{SE}}$, and $\mathrm{Random}_{\mathrm{SE}}$ denote degree-based, in-degree-based, betweenness-based, PageRank-based, SCT-based, FCT-based, and random removal sequences, respectively. Bold values indicate the top three values within each SE metric. Larger values indicate higher robustness. The average row represents the average secondary-extinction response of each sequence.

| Food web | D_SE | ID_SE | BET_SE | PR_SE | SCT_SE | FCT_SE | Random_SE |
|---|---|---|---|---|---|---|---|
| AkatoreB | 0.0345 | 0.0345 | 0.0345 | 0.0345 | 0.0345 | 0.0345 | 0.0345 |
| Blackrock | 0.0345 | 0.0345 | 0.0345 | 0.0345 | 0.0345 | 0.0345 | 0.0345 |
| Broad | 0.0316 | 0.0316 | 0.0316 | 0.0316 | 0.0211 | 0.0316 | 0.0316 |
| Catlins | 0.0204 | 0.0204 | 0.0204 | 0.0204 | 0.0204 | 0.0204 | 0.0204 |
| DempstersAu | 0.0233 | 0.0349 | 0.0233 | 0.0349 | **0.0349** | 0.0349 | 0.0349 |
| DempstersSp | 0.0309 | 0.0309 | 0.0206 | 0.0309 | 0.0309 | 0.0309 | 0.0309 |
| German | 0.0349 | **0.0465** | **0.0349** | **0.0465** | **0.0349** | 0.0349 | **0.0465** |
| Kyeburn | 0.0306 | 0.0306 | 0.0306 | 0.0306 | 0.0306 | 0.0306 | **0.0408** |
| LilKyeburn | **0.0385** | **0.0385** | **0.0385** | **0.0385** | **0.0385** | **0.0385** | **0.0385** |
| Martins | 0.0286 | 0.0286 | 0.0190 | 0.0286 | 0.0190 | 0.0190 | 0.0286 |
| NorthCol | 0.0256 | 0.0256 | 0.0256 | 0.0256 | 0.0128 | 0.0256 | 0.0256 |
| Powder | 0.0256 | 0.0256 | 0.0256 | 0.0256 | 0.0256 | 0.0256 | 0.0256 |
| Stony | **0.0354** | 0.0354 | 0.0265 | 0.0354 | 0.0265 | **0.0354** | 0.0354 |
| SuttonSp | **0.0380** | **0.0380** | **0.0380** | **0.0380** | 0.0253 | **0.0380** | 0.0380 |
| Troy | 0.0256 | 0.0256 | 0.0256 | 0.0256 | 0.0256 | 0.0256 | 0.0256 |
| Venlaw | 0.0290 | 0.0290 | 0.0290 | 0.0290 | 0.0290 | 0.0290 | 0.0290 |
| **Average** | 0.0304 | 0.0319 | 0.0286 | 0.0319 | 0.0278 | 0.0306 | 0.0325 |

#### Empirical food webs MVSS trophic-class composition

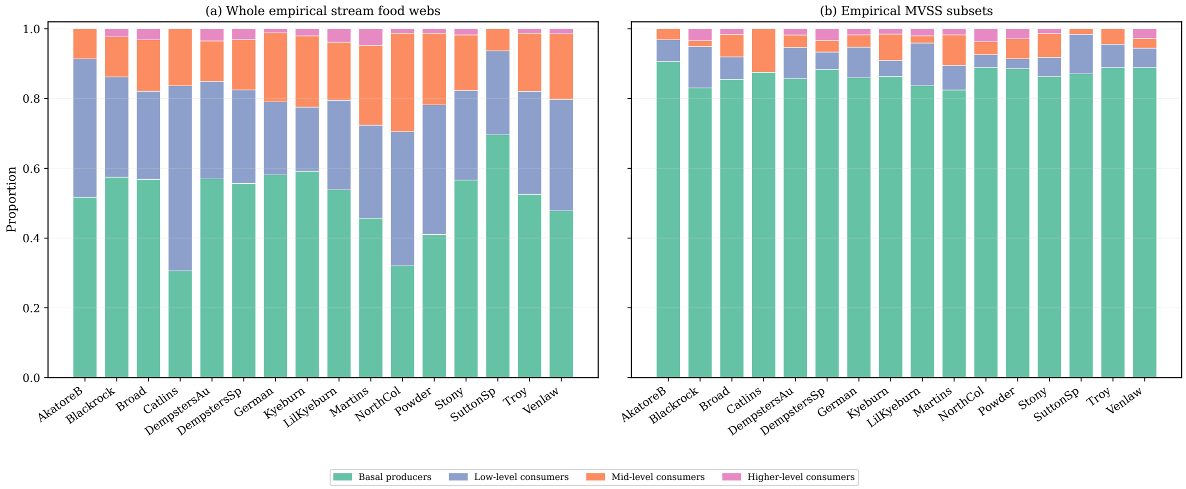


Figure S9: Broad trophic-class composition of the 16 empirical stream food webs and their MVSS subsets. Species trophic levels were calculated using the same prey-averaged trophic-level formula (following Levine (1980) and Williams and Martinez (2004)) used for the virtual food-web trophic-composition analysis and then grouped into broad trophic classes for display. Panel (a) shows the trophic-class composition of the whole empirical stream food webs, and panel (b) shows the corresponding composition of the MVSS subsets.
